# Overcoming Resistance to BRAF^V600E^ Inhibition in Melanoma by Deciphering and Targeting Personalized Protein Network Alterations

**DOI:** 10.1101/2020.11.03.366245

**Authors:** S. Vasudevan, E. Flashner-Abramson, I. Adesoji Adejumobi, D. Vilencki, S. Stefansky, A.M. Rubinstein, N. Kravchenko-Balasha

**Affiliations:** Department for Bio-medical Research, Faculty of Dental Medicine, Hebrew University of Jerusalem, Jerusalem 91120, Israel

**Keywords:** Melanoma, information theory, surprisal analysis, patient-specific altered signaling signatures, personalized therapy, BRAF^V600E^-mutated melanoma

## Abstract

BRAF^V600E^ melanoma patients, despite initially responding to the clinically prescribed anti-BRAF^V600E^ therapy, often relapse and their tumors develop drug resistance. While it is widely accepted that these tumors are originally driven by the BRAF^V600E^ mutation, they often eventually diverge and become supported by various signaling networks. Therefore, patient-specific altered signaling signatures should be deciphered and treated individually.

In this study, we design individualized melanoma combination treatments based on personalized network alterations. Using an information-theoretic approach, we compute high-resolution patient-specific altered signaling signatures. These altered signaling signatures each consist of several co-expressed subnetworks, which should all be targeted to optimally inhibit the entire altered signaling flux. Based on these data, we design smart, personalized drug combinations, often consisting of FDA-approved drugs. We validate our approach in vitro and in vivo showing that individualized drug combinations that are rationally based on patient-specific altered signaling signatures are more efficient than the clinically used anti-BRAF^V600E^ or BRAF^V600E^/MEK targeted therapy. Furthermore, these drug combinations are highly selective, as a drug combination efficient for one BRAF^V600E^ tumor is significantly less efficient for another, and vice versa. The approach presented herein can be broadly applicable to aid clinicians to rationally design patient-specific anti-melanoma drug combinations.

## Introduction

The rates of melanoma have been rapidly increasing [1]. Melanoma is one of the most common cancers in young adults, and the risk for melanoma increases with age [1]. However, alongside the rapid increase in incidence, there has also been rapid clinical advancement over the past decade, with targeted therapy and immunotherapy that have become available to melanoma patients [2].

Melanoma is associated with a great burden of somatic genetic alterations [3], with the primary actionable genomic data being an activating mutation in the BRAF gene, BRAF^V600E^, occurring in ∼50% of all melanomas [3,4].

Nearly a dozen new treatments have been approved by the Food and Drug Administration (FDA) for unresectable or metastatic melanoma harboring the BRAF^V600E^ mutation, among them vemurafenib (a BRAF^V600E^ inhibitor), cobimetinib (a MEK^MAPK^ inhibitor), or a combination of dabrafenib and trametinib (a BRAF^V600E^ inhibitor and a MEK^MAPK^ inhibitor, respectively) [2].

While targeted therapy revolutionized melanoma treatment, the high hopes shortly met a disappointment, as it became evident that most patients treated with BRAF^V600E^ inhibitors eventually relapse and their tumors become resistant to the treatment [5–7]. Various combination treatments were suggested to overcome the acquired resistance to BRAF^V600E^ inhibitors [5,6,9,10]. Nevertheless, BRAF^V600E^ and MEK inhibitors remain the only targeted agents approved by the FDA for melanoma.

In this study, we design patient-specific targeted treatments for melanoma based on individualized alterations in signaling protein networks, rather than on genomic or protein biomarkers. Attempting to treat patients based on the identification of single biomarkers or signaling pathways may overlook tumor-specific molecular alterations that have evolved during the course of disease, and the consequently selected therapeutic regimen may lack long term efficacy resulting from partial targeting of the tumor imbalance. We have shown that different patients may display similar oncogene expression levels, albeit carrying biologically distinct tumors that harbor different sets of unbalanced molecular processes [11]. Therefore, we suggest to explore the cancer data space utilizing an information theoretic approach that is based on surprisal analysis [11–13], to unbiasedly identify the altered signaling network structure that has emerged in every single tumor [11,12].

Our thermodynamic-like viewpoint grasps that tumors are altered biological entities, which deviate from their steady state due to patient-specific alterations. Those alterations can manifest in various manners that are dependent on environmental or genomic cues (e.g. carcinogens, altered cell-cell communication, mutations, etc.) and give rise to one or more distinct groups of co-expressed onco-proteins in each tumor, named unbalanced processes [11–13]. A patient-specific set of unbalanced processes constitutes a unique signaling signature and provides critical information regarding the elements in this signature that should be targeted. Each tumor can harbor several distinct unbalanced processes, and therefore all of them should be targeted in order to collapse the altered signaling flux in the tumor [11,12]. We have demonstrated that with comprehensive knowledge about the patient-specific altered signaling signature (PaSSS) in hand, we can predict highly efficacious personalized combinations of targeted drugs in breast cancer [12]. Herein, we decipher the accurate network structure of co-expressed functional proteins in melanoma tumors, hypothesizing that the PaSSS identified will guide us on how to improve the clinically used BRAF^V600E^-targeted drug combinations. Our aim was to examine the ability of PaSSS-based drug combinations to reduce the development of drug resistance, which frequently develops following BRAF^V600E^ inhibition in melanoma.

To this end, we studied a dataset consisting of 353 BRAF^V600E^ and BRAF^WT^ skin cutaneous melanoma (SKCM) samples, aiming to gain insights into the altered signaling signatures that have emerged in these tumors. A set of 372 thyroid carcinoma (THCA) samples was added to the dataset, as these tumors frequently harbor BRAF^V600E^ as well, therefore enabling studying the commonalities and differences between tumor types that frequently acquire the BRAF^V600E^ mutation.

We show that 17 distinct unbalanced processes are repetitive among the 725 SKCM and THCA patient-derived cancer tissues. Each tumor is characterized by a specific subset of typically 1-3 unbalanced processes. Interestingly, we demonstrate that the PaSSS does not necessarily correlate with the existence of the BRAF^V600E^, namely different tumors can harbor different signatures while both carrying the mutated BRAF, and vice versa – tumors can harbor the same altered signaling signature regardless of whether they carry BRAF^V600E^ or BRAF^WT^. These data suggest that examination of the BRAF gene alone does not suffice to tailor effective medicine to the patient. SKCM and THCA patients harboring BRAF^V600E^ can respond differently to the same therapeutic regimen, or rather benefit from the same treatment even though their BRAF mutation status differs.

We experimentally demonstrate our ability to predict effective personalized therapy by analyzing a cell line dataset and tailoring efficacious personalized combination treatments to two BRAF^V600E^-harboring melanoma cell lines, A375 and G361. The predicted PaSSS-based drug combinations were shown to have an efficacy superior to monotherapies or other drug combinations (such that were not predicted to target the individualized altered signaling signatures, and combinations used in clinics), both *in vitro* and *in vivo*. We show that an in depth resolution of individualized signaling signatures allows inhibiting the development of drug resistance and melanoma regrowth, by demonstrating that while A375 and G361 melanomas develop drug resistance several weeks following initial administration of the clinically used combination, dabrafenib+trametinib, individualized PaSSS-based drug combinations gain a longer lasting effect and show high selectivity.

## Methods

### Datasets

This study utilized a protein expression dataset consisting of 353 skin cutaneous melanoma (SKCM) sample and 372 thyroid carcinoma (THCA) samples. The samples were selected from a large TCPA dataset containing 7694 cancer tissues from various anatomical origins (PANCAN32, level 4 [14]). Each cancer tissue was profiled on a reverse phase protein array (RPPA) for 258 cancer-associated proteins. After filtering out proteins that had NA values for a significant number of patients, 216 proteins remained for further analysis.

The dataset for the cancer cell lines was downloaded from the TCPA portal [14]. The data was already published by Li et al. [15]. A part of the original dataset containing 290 cell lines from 16 types of cancers was selected, including breast, melanoma, ovarian, brain, blood, lung, colon, head and neck, kidney, liver, pancreas, bone and different types of sarcomas, stomach-oesophagus, uterus and thyroid cancers. The cell lines in the dataset were profiled for 224 phospho-proteins and total proteins using RPPA.

### Surprisal analysis

Surprisal analysis is a thermodynamic-based information-theoretic approach [16–18]. The analysis is based on the premise that biological systems reach a balanced state when the system is free of constraints [19–21]. However, when under the influence of environmental and genomic constraints, the system is prevented from reaching the state of minimal free energy, and instead reaches a state which is higher in free energy (in biological systems, which are normally under constant temperature and constant pressure, minimal free energy equals maximal entropy).

Surprisal analysis can take as input the expression levels of various macromolecules, e.g. genes, transcripts, or proteins. However, be it environmental or genomic alterations, it is the proteins that constitute the functional output in living systems, therefore we base our analysis on proteomic data. The varying forces, or constraints, that act upon living cells ultimately manifest as alterations in the cellular protein network. Each constraint induces a change in a specific part of the protein network in the cells. The subnetwork that is altered due to the specific constraint is termed an unbalanced process. System can be influenced by several constraints thus leading to the emergence of several unbalanced processes. When tumor systems are characterized, the specific set of unbalanced processes is what constitutes the tumor-specific signaling signature.

Surprisal analysis discovers the complete set of constraints operating on the system in any given tumor, *k*, by utilizing the following equation [22]: ln 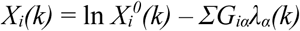, where *i* is the protein of interest, 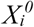 is the expected expression level of the protein when the system is at the steady state and free of constraints, and *ΣG*_*iα*_*λ*_*α*_*(k)* represents the sum of deviations in expression level of the protein *i* due to the various constraints, or unbalanced processes, that exist in the tumor *k*.

The term *G*_*α*_ denotes the degree of participation of the protein *i* in the unbalanced process α, and its sign indicates the correlation or anti-correlation between proteins in the same process (**Table S1**). Proteins with significant *G*_*iα*_ values are grouped into unbalanced processes (**Fig. S1, Table S2**) that are active in the dataset [12].

The term *λ*_*α*_*(k)* represents the importance of the unbalanced process α in the tumor *k* (**Table S1**). The partial deviations in expression level of the protein *i* due to the different constraints sum up to the total change in expression level (relative to the balance state level), *ΣG*_*iα*_*λ*_*α*_*(k)*.

For complete details regarding the analysis please refer to the SI of reference 12.

### Determination of the number of significant unbalanced processes

The analysis of the 725 patients provided a 725×216 matrix of *λ*_*α*_*(k)* values, such that every row in the matrix contained 216 values of *λ*_*α*_*(k)* for 725 patients, and each row corresponded to an unbalanced process (**Table S1**). However, not all unbalanced processes are significant. Our goal is to determine how many unbalanced processes are needed to reconstruct the experimental data, i.e. for which value of *n*: ln (*Xi(k)/M) ≈ - ΣG*_*iα*_*λ*_*α*_*(k)*. To find *n*, we performed the following two steps:

#### (1) Reproduction of the experimental data by the unbalanced processes was verified

We plotted *ΣG*_*iα*_*λ*_*α*_*(k)* for *α =1, 2,…, n* against ln *X*_*i*_*(k)* for different proteins, *i*, and for different values of *n*, and examined the correlation between them as *n* was increased. An unbalanced process, *α = n*, was considered significant if it improved the correlation significantly relative to *α = n – 1* (**Fig. S2**) (see [11] for more details).

#### (2) Processes with significant amplitudes were selected

To calculate threshold limits for *λ*_*α*_*(k)* values (presented in **Table S1** and **Fig. S3**) the standard deviations of the levels of the 10 most stable proteins in this dataset were calculated (e.g. those with the smallest standard deviations values). Those fluctuations were considered as baseline fluctuations in the population of the patients which are not influenced by the unbalanced processes. Using standard deviation values of these proteins the threshold limits were calculated as described previously [23]. The analysis revealed that from α = 18, the importance values, *λ*_*α*_*(k)*, become insignificant (i.e. do not exceed the noise threshold), suggesting that 17 unbalanced processes are enough to describe the system.

For more details see references 12 and 22.

### Generation of functional subnetworks

The functional sub-networks presented in Figures 2, 5, 6, S1 and S4 were generated using a python script as described previously [12]. Briefly, the goal was to generate a functional network according to STRING database, where proteins with negative G values are marked blue and proteins with positive G values are marked red, to easily identify the correlations and anti-correlations between the proteins in the network. The script takes as an input the names of the genes in the network and their G values, obtains the functional connections and their weights from STRING database (string-db.org), and then plots the functional network (using matplotlib library).

**Figure 2:**
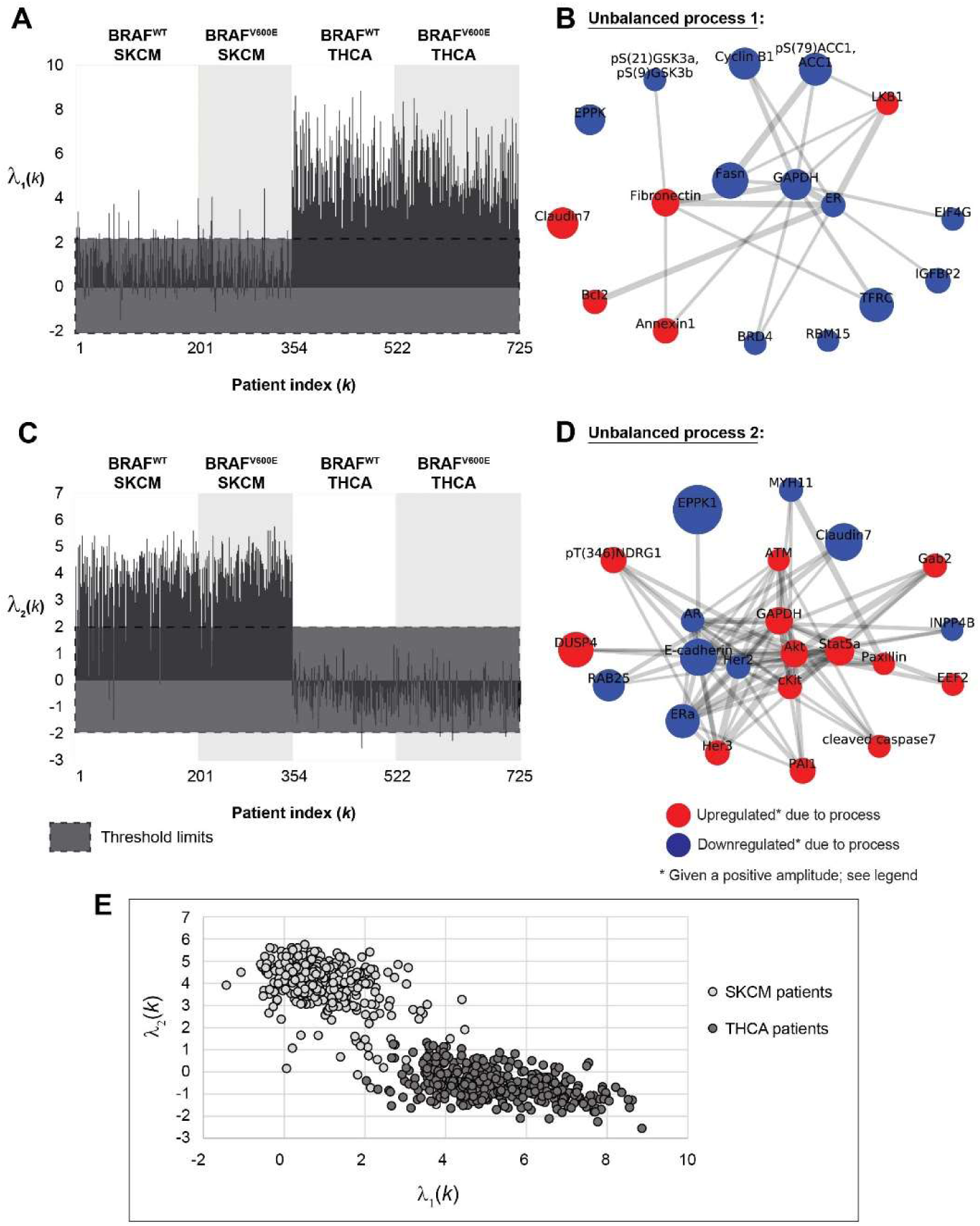
Unbalanced processes 1 and 2 distinguish well between SKCM and THCA tumors when plotted in 2D. The majority of THCA tumors harbor unbalanced process 1 (**A**), while the majority of SKCM tumors harbor unbalanced process 2 (**C**). Unbalanced processes 1 and 2 are shown in panels **B** and **D. <u>Note</u>** that red proteins are upregulated, and blue proteins are downregulated given that the amplitude of the process is positive. In tumors where the amplitude is negative, the direction of change is opposite. (**E**) A 2D plot showing λ_2_(k) against λ_1_(k) for all SKCM and THCA patients. The plot shows nicely the separation between SKCM and THCA patients in this 2D space. Note, however, that every tumor is characterized by a set of unbalanced processes (a PaSSS), and that unbalanced processes 1 and 2 alone do not suffice to describe the complete tumor-specific altered signaling signatures.

**Figure 5:**
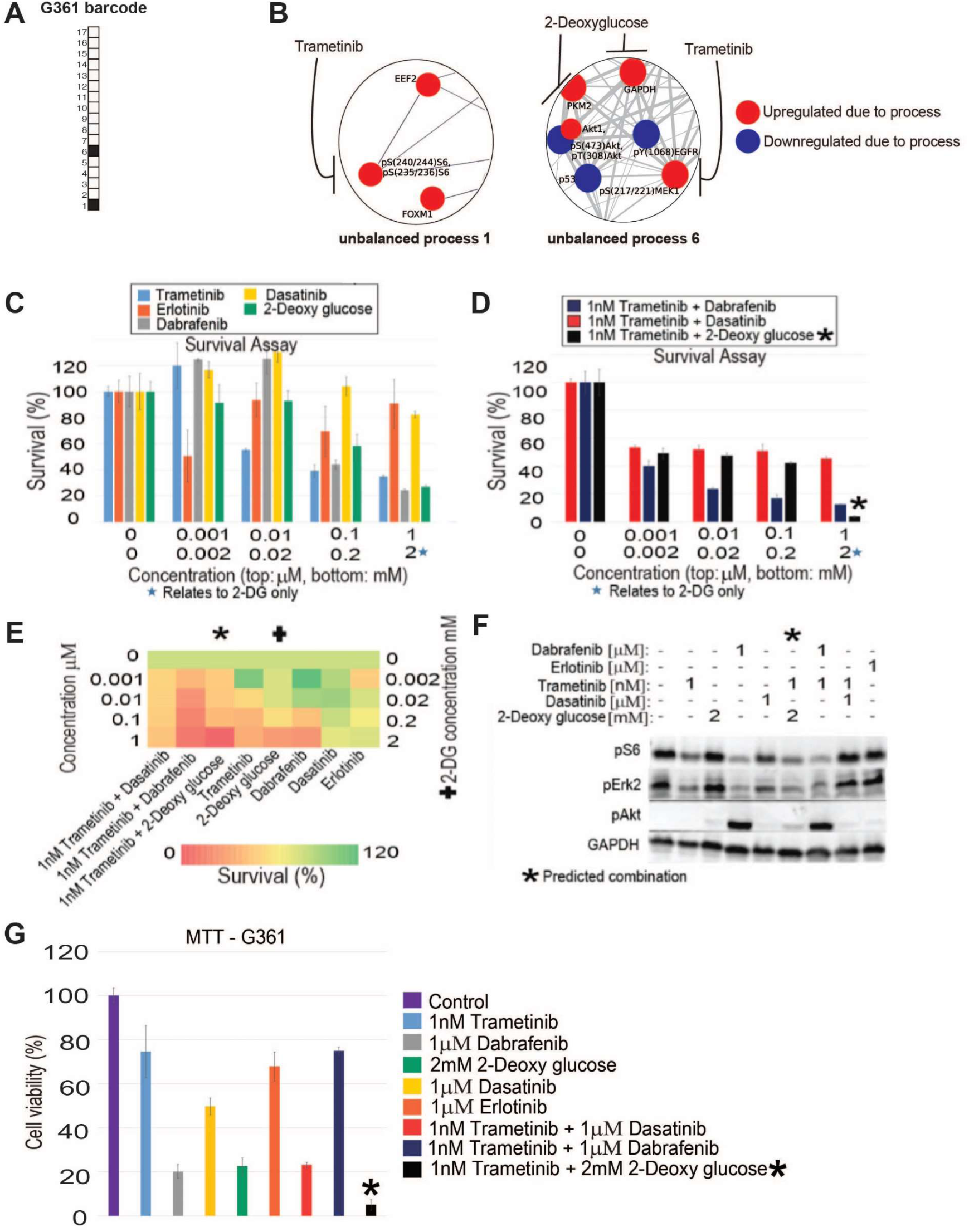
G361 melanoma cells altered signaling signature and treatment. (**A**) Barcode of active unbalanced processes for G361 based on PaSSS analysis. (**B**) Zoom in images of the active unbalanced processes, 1 and 6, in G361 cells, as well as the drugs targeting the central proteins in each unbalanced process. The upregulated proteins are colored red and the down regulated proteins are colored blue. (**C, D**) Survival rates of cells in response to different therapies. The cells were treated with the predicted combination (*) to target G361, the treatments used in the clinics for BRAF mutated melanoma malignancies, monotherapies of each treatment and the predicted combination used to target BRAF mutated melanoma cell line A375. The combination predicted to target G361 was more efficient than any other treatment. (**E**) Results of the survival assay (shown in panels A and B) are shown as a heatmap. (**F**) Western blot results after treatment with different therapies. The predicted combination depletes the signaling in G361 cells as represented by decrease in phosphorylation levels of pS6, pERK and pAkt. Akt remains active when the cells are treated with dabrafenib or dabrafenib + trametinib. (**G**) G361 cells were treated as indicated for 72 hours and then the viability of the cells was measured in an MTT assay. The effect of the predicted combination (marked in the with an asterix sign) was superior to combinations and single drugs expected to partially inhibit the cell line-specific altered signaling signature.

**Figure 6:**
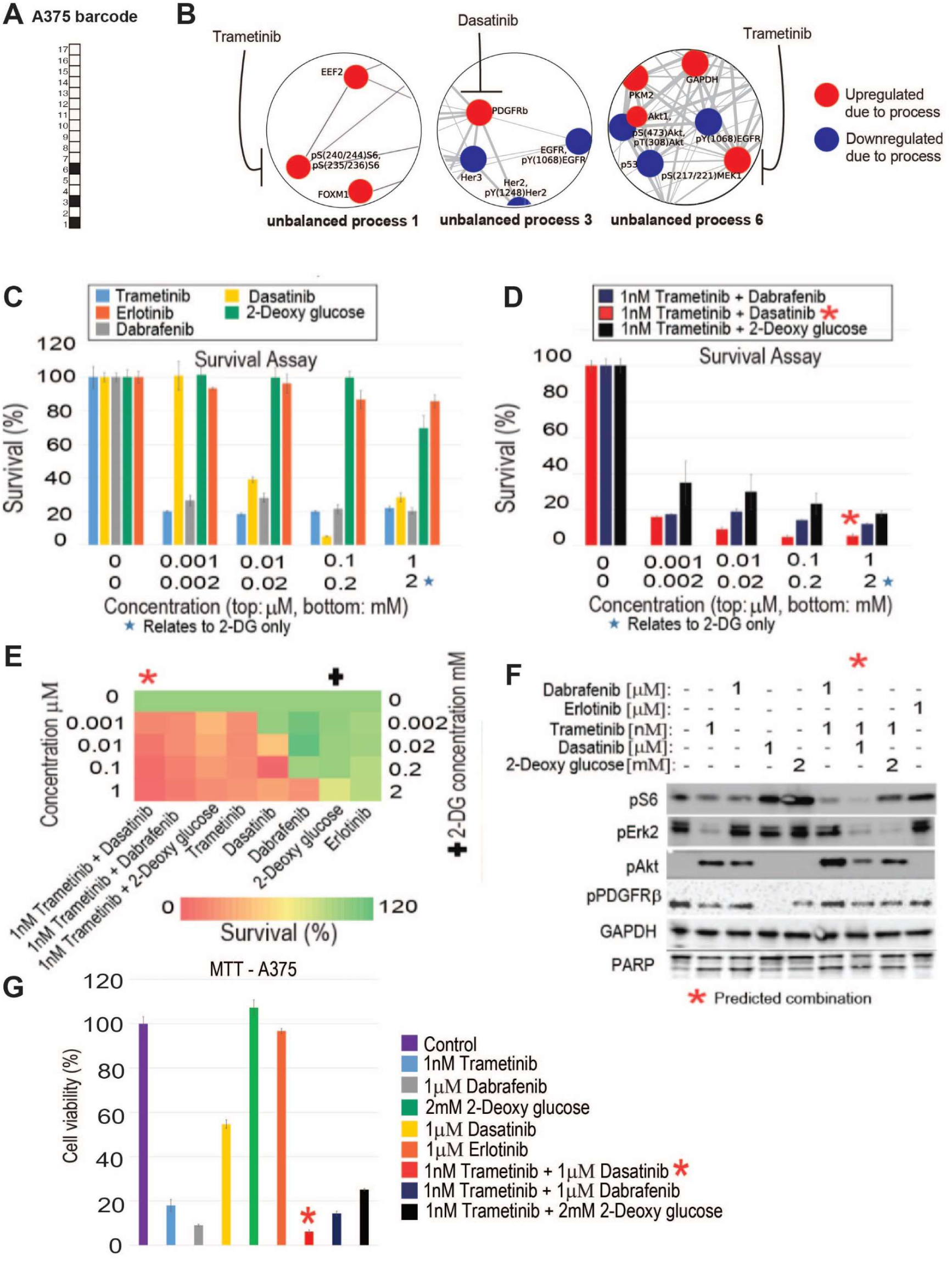
A375 melanoma cells altered signaling signature and SA-based treatment. Even though A375 cells harbor BRAF^V600E^, as do G361 cells, they were found to be characterized by a different set of active unbalanced processes, or PaSSS. (**A**) Barcode of the unbalanced processes for A375 based on PaSSS analysis. (**B**) Zoom in images of the active unbalanced processes, 1, 3 and 6, in A375 cells, as well as the drugs targeting the central proteins in each unbalanced process. The upregulated proteins are colored red and the down regulated proteins are colored blue. (**C, D**) Survival rates of cells in response to different therapies. The cells were treated with the predicted combination (*) to target A375, the treatments used in the clinics for BRAF mutated melanoma malignancies, monotherapies of each treatment and the predicted combination used to target BRAF mutated melanoma cell line G361. The combination predicted to target A375 was more efficient than any other treatment. (**E**) Results of the survival assay (shown in panels A and B) are shown as a heatmap. (**F**) Western blot results after treatment with different therapies. The predicted combination depletes the signaling in A375 cells as represented by decrease in phosphorylation levels of pS6, pERK, pAkt and pPDGFRβ. Akt remains active when the cells are treated with monotherapies - trametinib or dabrafenib, and the combination therapies - dabrafenib + trametinib or trametinib + 2-deoxy glucose, the predicted combination of G361. (**G**) A375 cells were treated as indicated for 72 hours and then the viability of the cells was measured in an MTT assay. The effect of the predicted combinations (marked in the figure with asterix signs) was superior to combinations and single drugs expected to partially inhibit the cell line-specific altered signaling signature.

### Barcode calculation

The barcodes of unbalanced processes were generated using a python script. For each patient, *λ*_*α*_*(k)* (α = 1, 2, 3,…, 17) values were normalized as follows: If *λ*_*α*_*(k) > 2* (and is therefore significant according to calculation of threshold values) then is was normalized to 1; if *λ*_*α*_*(k) < −2* (significant according to threshold values as well) then it was normalized to −1; and if *−2 < λ*_*α*_*(k) < 2* then it was normalized to 0.

### Cell Culture

The BRAF mutated melanoma cell lines, A375 and G361, were obtained from the ATCC and grown in DMEM (G361) or RPMI (A375) medium. The cells were supplemented with 10 % fetal calf serum (FCS), L-glutamine (2mM), 100 U/ml penicillin and 100 mg/ml streptomycin and incubated at 37 °C in 5% CO_2_. The cell lines were authenticated at the Biomedical Core Facility of the Technion, Haifa, Israel.

### Western blot analysis

The cells were seeded into 6 well plates (∼1.5 = 10^6^ cells/well) and grown under complete growth media. A375 cells were treated the next day as indicated for 48 hours in partial starvation medium (RPMI medium with 1.2% FCS). G361 cells were treated in complete growth medium for 24 hours. The dead cells were collected from the medium. The adherent cells were then treated with IGF for 15 minutes. The cells were then lysed using hot sample buffer (10% glycerol, 50 mmol/L Tris-HCl pH 6.8, 2% SDS, and 5% 2-mercaptoethanol) and western blot analysis was carried out. The lysates were fractionated by SDS-PAGE and transferred to nitrocellulose membranes using a transfer apparatus according to the manufacturer’s protocols (Bio-Rad). Blots were developed with an ECL system according to the manufacturer’s protocols (Bio-Rad).

### Methylene blue assay

In a 96 well plate, the cells were seeded and treated as indicated for 72 hours. The cells were fixed with 4% paraformaldehyde and then stained with methylene blue. To calculate the number of surviving cells, the color was extracted by adding 0.1M Hydrochloric acid and the absorbance was read at 630 nm.

### MTT assay

Cells were seeded and treated as indicated in a 96 well plate for 72 hours. Cell viability was checked using MTT assay kit (Abcam). Equal volumes of MTT solution and culture media were added to each well and incubated for 3 hours at 37 °C. MTT solvent was added to each well, and then the plate was covered in aluminum foil and put on the orbital shaker for 15 minutes. Absorbance was read at 590nm following 1 hour.

### Resistance Assay

Cells were seeded in multiple 96 well plates and treated as needed in various time points (3, 7, 14, 21, 28, 35, 42, 49, 54 days). At every time point the cells were fixed with 4% paraformaldehyde and then stained with methylene blue. The number of cells which survived at each time point was quantified by adding 0.1M Hydrochloric acid and reading the absorbance at 630 nm.

### Animal Studies

The cells - A375 (0.25 × 10^6^ cells/mouse) or G361 (0.5 × 10^6^ cells/ mouse) - were inoculated subcutaneously into NSG mice (n = 8 mice per group), and once the volume of the tumors reached 50 mm^3^, treatments were initiated 6 times a week for up to 4 weeks. Tumor volume was measured twice a week. Trametinib (0.5mg/kg), dasatinib (35mg/kg) and dabrafenib (35mg/kg) were suspended in an aqueous mixture of 0.5% hydroxypropyl methylcellulose + 0.2% tween 80 and administered by oral gavage. 2-deoxy-D-glucose (500mg/kg) was suspended in saline and injected intraperitoneally. All the drugs were purchased from Cayman chemicals (Enco, Israel). The Hebrew University is an AAALAC International accredited institute. All experiments were conducted with approval from the Hebrew University Animal Care and Use Committee. Ethical accreditation number: Md-17-15174-4.

## Results

### An overview of the experimental-computational approach

Biomarker analysis in melanoma relies mainly on the identification of mutations in the BRAF gene [24]. If mutation/upregulation of the mutant BRAF^V600E^ is identified (**Fig. 1**, left), the patient will likely be treated with a BRAF^V600E^ inhibitor (e.g. vemurafenib [25] or dabrafenib [26]), possibly concurrently with an inhibitor of MEK^MAPK^ (e.g. trametinib [27]). The combination of BRAF^V600E^ and MEK^MAPK^ inhibitors was shown to be superior to BRAF^V600E^ inhibition alone and to delay or prevent the development of drug resistance [9].

**Figure 1:**
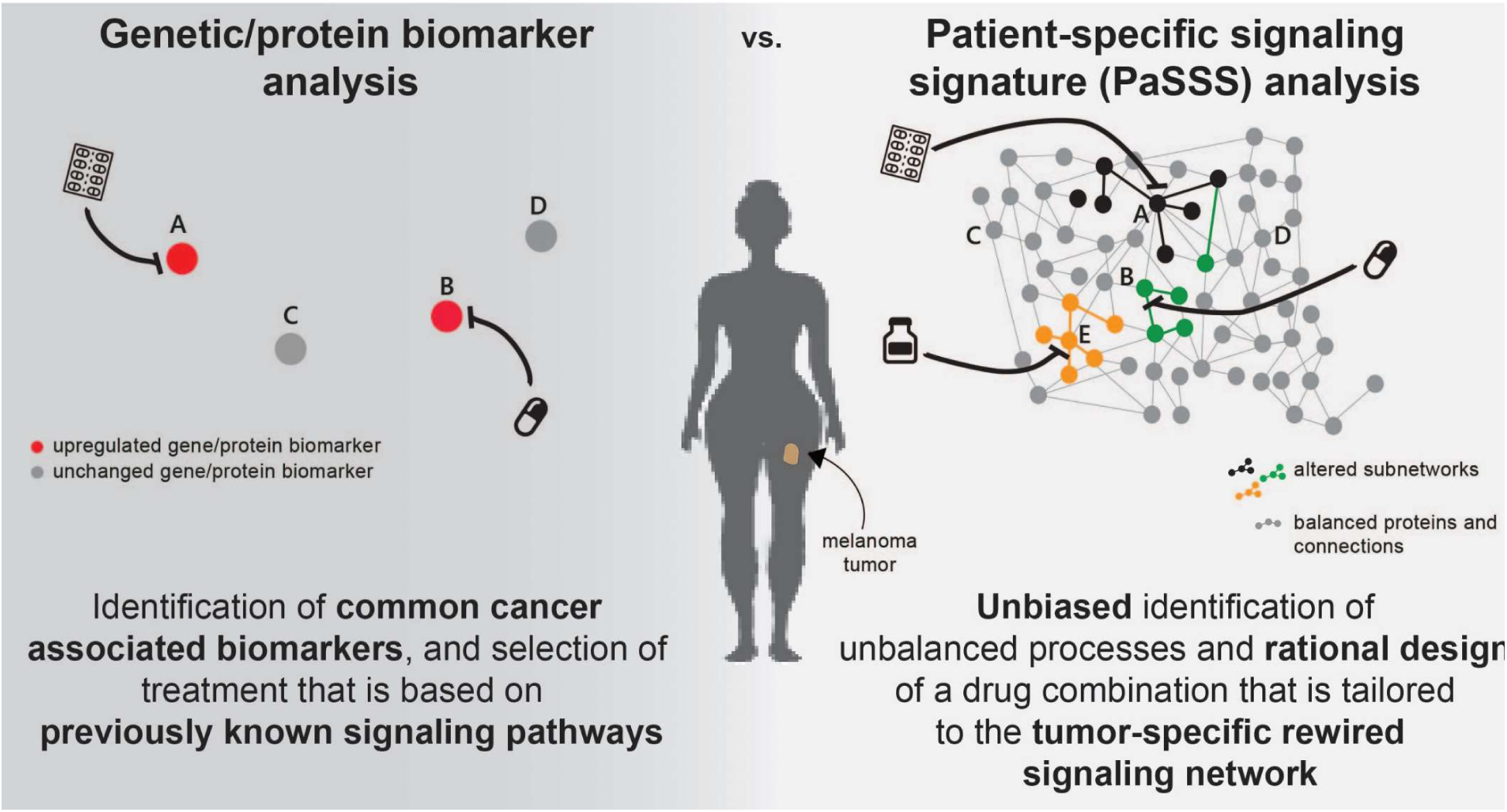
Conventional biomarker analysis vs. patient-specific signaling signature analysis. Genetic/protein biomarker analysis relies on evaluation of the expression levels of common cancer type-associated genes or proteins (left). The design of a drug combination is done according to inference of the state of the surrounding signaling network, based on previous knowledge (left). In contrast, patient-specific signaling signature (PaSSS) analysis involves proteomic analysis of hundreds of cancer-associated proteins, and unbiased identification of the altered signaling signature in every sample, i.e. that does not depend on previous knowledge of signaling pathways. This enables rationally designing personalized combinations of targeted drugs that are based on the patient-specific uniquely rewired signaling network (right).

However, the biomarker analysis utilized in clinics lacks information about the altered signaling network, and, for example, may overlook additional or alternative protein targets that, if targeted by drugs, may enhance the efficacy of the treatment (**Fig. 1**, left).

We utilize an information theoretic approach that is based on surprisal analysis (**Methods**) [11–13] to gain information regarding the patient-specific signaling signature (PaSSS) that has emerged in every individual tumor (**Fig. 1**, right). Based on proteomic analysis of the samples, we identify the set of altered protein-protein co-expressed subnetworks, or *unbalanced signaling processes*, that has arisen as a result of constraints (environmental or genomic) which operate on the tumor, and then design a combination of targeted drugs that is expected to collapse the tumor-specific altered signaling signature (**Fig 1**, right and **Methods**) [11–13].

We obtained from the TCPA database [14] a dataset containing 353 skin cutaneous melanoma (SKCM) and 372 thyroid cancer (THCA) samples (725 samples in total). The thyroid cancer samples were added to the dataset for two main reasons: (1) to increase the number of samples in the dataset, thereby increasing the resolution of the analysis; (2) THCA tumors frequently harbor the BRAF^V600E^ mutation, and we were therefore interested in examining the commonalities and differences between the altered signaling signatures that emerged in SKCM and THCA tumors.

### 17 unbalanced processes repeat themselves throughout 725 SKCM and THCA tumors

The analysis of the dataset revealed that the 725 SKCM and THCA tumors can be described by 17 unbalanced processes (**Fig. S1**; the amplitudes for each process in each patient and the importance of each protein in the different processes can be found in **Table S1**; the protein composition of each process is presented in **Table S2**), i.e. 17 distinct unbalanced processes suffice to reproduce the experimental data (**Fig. S2** and **Methods**).

Unbalanced processes 1 and 2, the two most significant unbalanced processes, which appear in the largest number of tumors, distinguish well between SKCM and THCA tumors, as can be seen by the 2D plots of λ_α_(k) values (i.e. amplitudes of each process in every tumor; **Fig. 2A**,**C**,**E**). Unbalanced process 1 appears almost exclusively in THCA tumors (372 THCA tumors harbor unbalanced process 1, vs. 46 SKCM tumors; **Fig. 2A**,**E, Table S2**), while unbalanced process 2 characterizes almost exclusively SKCM tumors (331 SKCM tumors harbor unbalanced process 2, vs. only 4 THCA tumors; **Fig. 2C**,**E, Table S2**). Unbalanced process 1 involves upregulation of proteins that have been previously linked to THCA: LKB1 [28], fibronectin [29,30], Bcl-2 [31], claudin 7 [32] (**Fig. 2B**). Unbalanced process 2 is characterized by the upregulation of proteins that have been implicated in melanoma, such as Stat5α [33], Akt [34], cKit [35], Her3 [36], and ATM [37] (**Fig. 2D**). As can be seen in the graph in Figure 2C, unbalanced process 2 was assigned a positive amplitude in all 331 SKCM tumors in which it appears, while in 4 THCA tumors it was assigned a negative amplitude (see also **Table S2**). This means that the proteins that participate in this unbalanced process deviate to opposite directions in the two types of tumors (importantly, this remark denotes only the partial deviation that occurred in these proteins due to unbalanced process 2; some of these proteins may have undergone additional deviations due to the activity of other unbalanced processes. See **Table S2** and **Methods**). Although this dominant process appears in a significant number of BRAF^V600E^ SKCM patients (**Fig. 2C**), it does not include pS(445)BRAF and downstream signaling. This finding corresponds to a recent characterization of melanoma tissues [4] and suggests that the signaling signatures of BRAF^V600E^ tissues may diverge over time and acquire additional signaling routs which are not necessarily related to the original driver mutations, such as BRAF^V600E^ or its downstream MEK^MAPK^ signaling.

Unbalanced process 2 can also be found in BRAF^WT^ patients (**Fig. 2C**). See, for example, patient TCGA-XV-AAZV (**Fig. 3**). The signature of this patient did not include additional processes. A total of 181 SKCM patients harbor this signaling signature, consisting only of unbalanced process 2: 100 of them harbor BRAF^WT^ and 81 of them harbor BRAF^V600E^ (**Fig. 3**). In contrast, no THCA patients harbor this signature (**Fig. 3**). The finding that BRAF^WT^ and BRAF^V600E^ SKCM patients can, in some cases, harbor the same altered signature suggests that these patients can also benefit from the same combination of targeted drugs.

**Figure 3:**
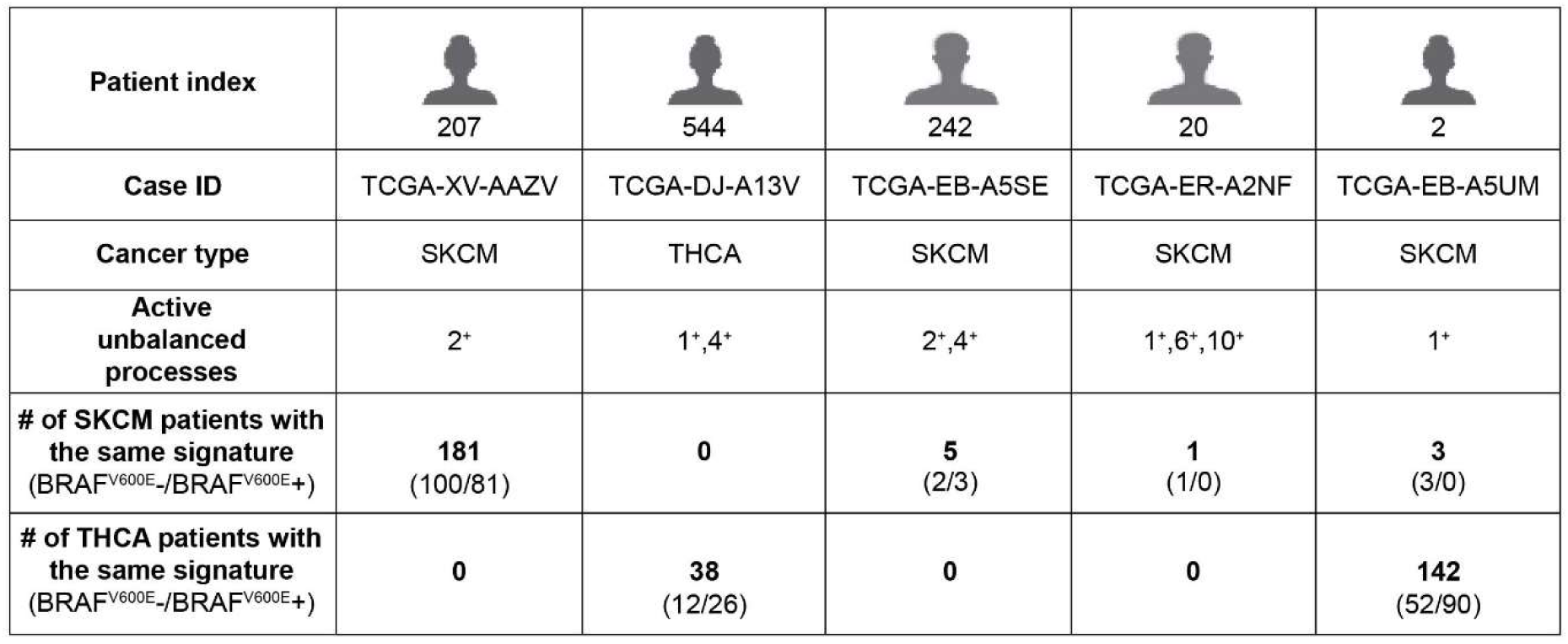
Examples for patient-specific sets of active unbalanced processes. Each patient typically harbors a set of 1-3 active unbalanced processes. Our results show that a specific *set* of active processes does not necessarily distinguish between BRAF^V600E^- and BRAF^V600E^+ patients, or between SKCM and THCA patients.

Although unbalanced processes 1 and 2 distinguish well between SKCM and THCA patients (**Fig. 2A**,**C**,**E**), these processes alone do not suffice to describe the PaSSS of all patients. Our analysis suggests that to decipher the altered signaling signature in every patient, 17 unbalanced processes should be considered. Hence, 2D plots may overlook important therapeutic information. When we inspect the patients in the context of a 17-dimensional space, where each dimension represents an unbalanced process, we find that not all SKCM patients harbor unbalanced process 2, and that those who do harbor this process may harbor additional unbalanced processes as well (**Fig. 3** and **Fig. S3**). We have shown that mapping the patients into a multi-dimensional space, a 17D space in our case, allows deciphering the set of unbalanced process, namely the PaSSS, in every tumor. This mapping is crucial for the design of efficacious treatments [12].

The SKCM patient TCGA-EB-A5SE, for example, is characterized by a PaSSS consisting of unbalanced processes 2 and 4 (**Fig. 3**). Only 5 SKCM patients were found to be characterized by this set of unbalanced processes. Two of the patients harbor BRAF^WT^ tumors, and 3 of them BRAF^V600E^ (**Fig. 3**).

The SKCM patient TCGA-ER-A2NF was found to harbor a PaSSS consisting of unbalanced processes 1, 6 and 10 (**Fig. 3**). This patient harbors a one-of-a-kind tumor, as no other patients in the dataset harbor this altered signaling signature (**Fig. 3**).

The PaSSS of THCA patient TCGA-DJ-A13V includes unbalanced processes 1 and 4 (**Fig. 3**). This signature characterizes 38 THCA patients, 12 of them BRAF^WT^ and 26 of them BRAF^V600E^ (**Fig. 3**). These THCA patients may benefit from a combination of drugs that target central protein nodes in unbalanced processes 1 and 4, regardless of whether they harbor BRAF^V600E^ or not. No SKCM patients harbor this altered signaling signature (**Fig. 3**).

Another interesting finding is that SKCM and THCA patients may harbor the same PaSSS, as is the case of the signature consisting of unbalanced process 1, shared by 3 SKCM patients and 142 THCA patients (**Fig. 3** and **Table S2**). All these patients may be treated with the same drug combination, targeting key proteins in unbalanced process 1, e.g. LKB1 and fibronectin (**Fig. 2B**).

### The altered signaling signatures identified in SKCM and THCA are almost mutually exclusive

To explore the entire dataset in terms of the set of unbalanced processes that each patient harbors, we assigned to each patient a patient-specific barcode, denoting the normalized PaSSS, i.e. the active unbalanced processes in the specific tumor, and the signs of their amplitudes (positive/negative) disregarding the size of the amplitude (**Fig. 4, Table S3**). These barcodes represent the mapping of every patient to a 17-dimensional space where each dimension denotes a specific unbalanced process [11,12]. We found that 138 distinct barcodes repeated themselves in the dataset (**Table S4**). Interestingly, the barcodes are almost mutually exclusive: 87 of the barcodes characterize SKCM tumors; 84 of them characterize <u>only</u> SKCM tumors and are not harbored by any THCA tumor (**Table S4**). 51 barcodes characterize THCA tumors; of them 48 characterize <u>solely</u> THCA tumors (**Table S4**).

**Figure 4:**
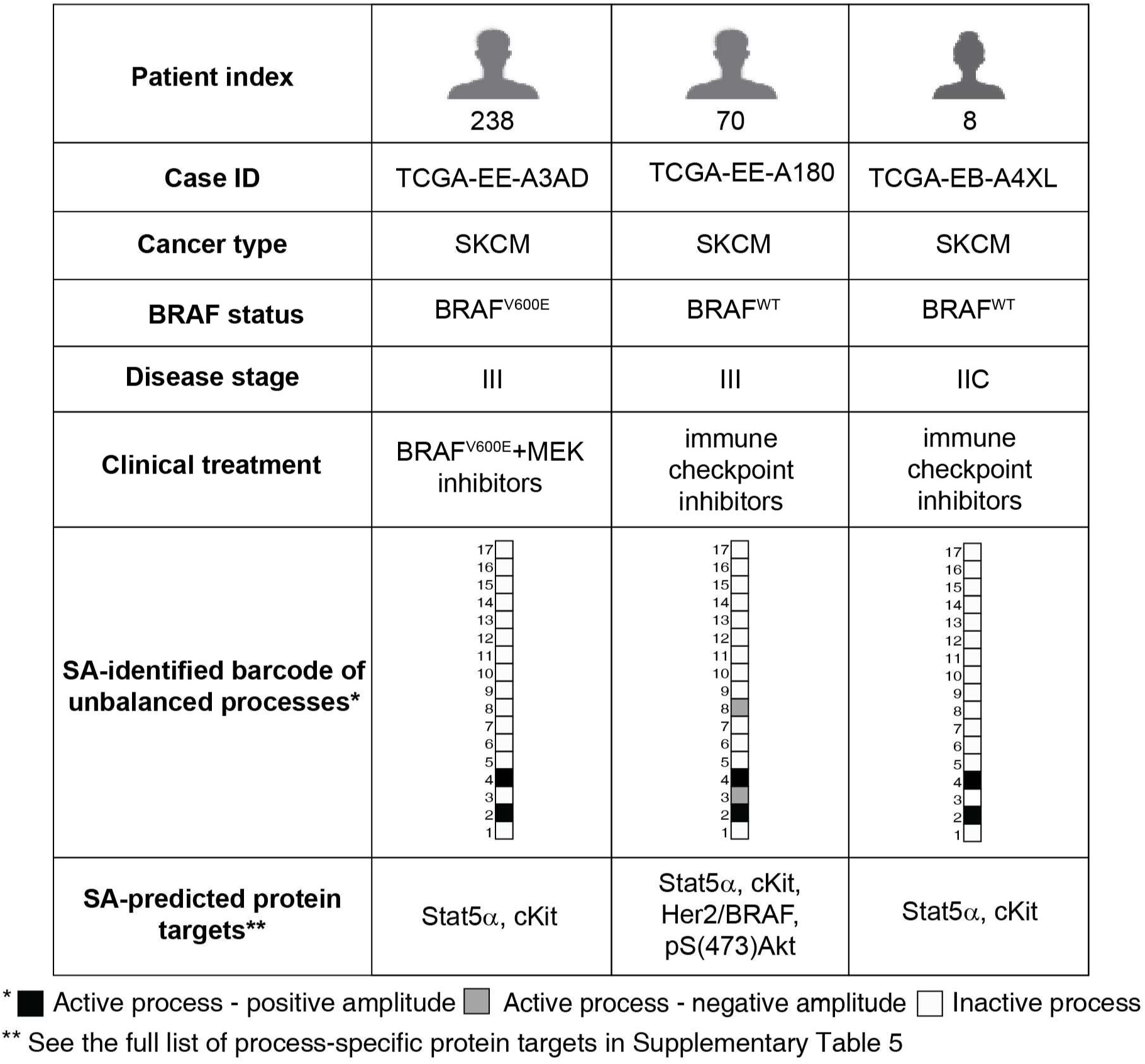
Patient-specific altered signaling signatures, or barcodes, can guide the design of personalized combination therapies. For each tumor, processes with amplitudes exceeding the threshold values (**Methods**) were selected and included in patient-specific sets of unbalanced processes. Those sets were converted into schematic barcodes. Central proteins from each process were suggested as potential targets for personalized drug combinations.

Most of the barcodes are rare: 81 barcodes are shared by only 5 SKCM tumors or less; 56 of them describe single, one-of-a-kind SKCM tumors (**Table S4**). 47 barcodes are shared by only 5 THCA tumors or less; 36 of them describe single THCA tumors (**Table S4**). This finding corroborates with our previous studies of signaling signatures in cancer [12], and underscores the need for personalized cancer diagnosis that is not biased by, e.g., the anatomical origin of the tumor.

### Patient-specific barcodes guide the rational design of personalized targeted combination therapy

We have previously shown the predictive power of our analysis in determining effective patient-tailored combinations of drugs that target key proteins in every unbalanced process [12,13].

Utilizing the maps of the unbalanced processes identified in the dataset herein (**Fig. S1**), we predicted process-specific protein targets for each process (**Table S5**). Each individual patient is predicted to benefit from a therapy that combines drugs against all the unbalanced processes active in the specific tumor (**Fig. 4, Table S5**).

As mentioned above, SKCM patients can in some cases benefit from the same combination therapy, regardless of their BRAF mutational status. This is the case for patients TCGA-EE-A3AD (carrying BRAF^V600E^) and TCGA-EB-A4XL (carrying BRAF^WT^), that were found to harbor tumors characterized by the same barcode of unbalanced processes, and were therefore predicted to benefit from the same treatment, where Stat5α and cKit are targeted simultaneously (**Fig. 4**).

Patient TCGA-EE-A180 carries BRAF^WT^, as does patient TCGA-EB-A4XL (**Fig. 4**). However, patient TCGA-EE-A180 harbors two active unbalanced processes that are not active in the tumor of patient TCGA-EB-A4XL – processes 3 and 8 (**Fig. 4**). Therefore, the list of proteins that should be targeted in order to collapse the tumor differs in these patients (**Fig. 4**).

### A375 and G361 melanoma cell lines harbor distinct altered signaling signatures

To experimentally validate our hypothesis that BRAF^V600E^ harboring cells may benefit from drug combinations that are designed based on the PaSSS identified at the time of diagnosis, we turned to analyze a different dataset containing 290 cell lines originating from 16 types of cancer, including blood, bone, breast, colon, skin, uterus, and more (see **Methods**). The cell lines were each profiled for the expression levels of 224 proteins and phosphoproteins using reverse phase protein assay.

PaSSS analysis of this cell line dataset revealed that 17 unbalanced processes were repetitive in the 291 cell lines (**Table S6, Table S7, Fig. S4**).

We selected two melanoma cell lines for experimental validation, G361 and A375. Both cell lines harbor the mutated BRAF^V600E^. In the clinic, patients bearing tumors with BRAF^V600E^ would all be treated similarly, with BRAF inhibitors alone or in combination with MEK inhibitors [9,27].

Our analysis, however, shows that G361 and A375 each harbor a distinct PaSSS (**Fig. 5**,**6**). G361 was found to harbor a PaSSS consisting of unbalanced processes 1 and 6 (**Fig. 5A**). The PaSSS of A375, on the other hand, consisted of three unbalanced processes, 1, 3 and 6 (**Fig. 6A**). The full lists of proteins participating in these processes are presented in Table S7, and images of the complete unbalanced processes can be found in Figure S4.

To predict cell line-specific drug combinations, central protein targets were selected from each active unbalanced process. In unbalanced process 6, pMEK1/2, GAPDH and PKM2 represent central upregulated proteins (**Fig. 5B, Fig. 6B**). Unbalanced process 3 was characterized by an upregulation of PDGFRβ (**Fig. 6B**), and unbalanced process 1 involved an upregulation of pS6 (**Fig. 5B, Fig. 6B**). We hypothesized that targeting these central proteins will reduce the signaling imbalance in A375 and G361 cell lines. Therefore, we predicted that G361 cells will be effectively treated by a drug combination containing trametinib (a pMEK1/2 inhibitor, commonly used for melanoma in clinics; also inhibits pS6 [38,39]) and 2-deoxy-D-glucose (2-DG; a glycolysis inhibitor, therefore affecting GAPDH and PKM2 levels; **Fig. 5B**). Based on the PaSSS of G361, trametinib should effectively target both unbalanced processes, 1 and 6 (**Fig. 5B**). However, since unbalanced process 6 was assigned a relatively high amplitude in G361 cells (**Table S6**), we predicted that a combination treatment combining trametinib with 2DG, which targets additional nodes in unbalanced process 6 (**Fig. 5B**), will more effectively target the PaSSS in G361 cells.

For A375, the amplitude of unbalanced process 6 was ∼2-fold lower than in G361 cells, and therefore we assumed that trametinib alone should suffice to efficiently reduce the signaling flux through this process. Thus, we predicted that a combination of trametinib and dasatinib (a multi-kinase inhibitor targeting also PDGFRβ) should effectively target the 3 unbalanced processes that constitute the PaSSS of these cells (**Fig. 6B**).

### The predicted drug combinations are cell line-specific and highly efficacious

Trametinib and dabrafenib, two clinically used drugs, indeed demonstrated relatively efficient killing of G361 cells, achieving up to ∼55% and ∼75% killing, respectively, when administered to the cells as monotherapies in a range of concentrations between 1 nM and 1 µM (**Fig. 5C**). Based on our analysis, we predicted that these drugs would each partially target the PaSSS in G361 cells (**Fig. 5A**,**B**).

Monotherapies of erlotinib and dasatinib were used as negative controls, as both were not expected to target the altered signaling signature of G361 cells (**Fig. 5C**,**D**). Indeed, both drugs demonstrated a weak effect on G361 cells, reaching up to ∼10% and ∼20% killing, respectively (**Fig. 5C**,**E**). 2-DG, which was predicted to target one of the unbalanced processes active in G361 cells, killed up to ∼70% of the cells at 2 mM (**Fig. 5C**,**E**).

When we tested combinations of drugs, we found that when G361 cells were treated with a combination of trametinib and dabrafenib, the clinically used combined treatment for BRAF^V600E^ melanoma, the combination was superior to each drug administered alone, and reached ∼90% killing of the cells when both drugs were administered at 1 µM (**Fig. 5D**,**E**). The results of our analysis, however, denoted that other major signaling nodes were altered in G361 cells, and that their targeting by drugs may be beneficial in these cells. When we tested the combination of trametinib and 2-DG, predicted by us to more effectively collapse the PaSSS that emerged in G361 cells, we indeed found that the combination abolished the cells almost completely when trametinib and 2-DG were added at 1 µM and 2mM, respectively (**Fig. 5D**,**E**). The combination of trametinib and 2-DG also effectively turned off the cellular signaling, as represented by the central proteins S6, Akt and ERK, while the other combinations we tested failed to do so (**Fig. 5F**). For example, the clinically used combination, dabrafenib+trametinib, induced the activity of pS(473)Akt (**Fig. 5F**), possibly reflecting a response of the cells to incomplete inhibition of the altered signaling flux.

In A375 cells, trametinib, dabrafenib and dasatinib killed up to ∼80% of the cells, when administered as single drugs (**Fig. 6C**,**E**). Erlotinib was used as a negative control, as it was predicted not to target any major node in the PaSSS of A375 cells, and indeed killed only up to ∼15% of the cells (**Fig. 6C**,**E**). 2-DG, which was predicted to target one of the three unbalanced processes active in A375 cells (**Fig. 6A**,**B**), killed up to ∼30% of the cells when administered as monotherapy (**Fig. 6C**,**E**). The clinically used drug combination, trametinib and dabrafenib, was more effective than each drug alone, and killed up to 90% of the cells (**Fig. 6D**,**E**). However, as in the case of G361 cells, we predicted that the clinically used combination would not be optimal in A375 cells, because another major node, PDGFRβ, should be targeted as well in order to effectively collapse the PaSSS in A375 cells (**Fig. 6A**,**B**). We therefore predicted that a combination of trametinib and dasatinib would efficiently target the altered signaling flux generated by 3 unbalanced processes in A375 cells (**Fig. 6A**,**B**). When we tested this combination, we found that it was highly efficacious and killed up to ∼95% of the cells (**Fig. 6D**,**E**). Moreover, trametinib and dasatinib, when combined, diminished S6 and ERK signaling, and lowered the levels of pPDGFRβ (**Fig. 6F**). As we found in the case of G361 cells, the clinically used combination, trametinib and dabrafenib, invoked an upregulation of pS(473)Akt in A375 cells as well (**Fig. 6F**). We tested the effect of combination predicted for G361 cells, trametinib and 2-DG, on A375 cells, and found that it was less effective in inhibiting the intra-cellular signaling (**Fig. 6F**) as well as cell survival than the drug combination predicted for the PaSSS of A375 (**Fig. 6D**,**E**). We attribute this finding to the fact the combination of trametinib and 2-DG targets only unbalanced processes 1 and 6, while leaving unbalanced process 3 untargeted, and therefore a partial effect is achieved by these drugs in A375 cells (**Fig. 6A**,**B**).

When tested in an MTT assay (assessing metabolic activity of the cells), the predicted combinations demonstrated higher efficacy and selectivity and were superior to other drug combinations or to each inhibitor alone (**Fig. 5G, 6G**).

### As opposed to common therapies used in clinics, the rationally designed cell line-specific drug combinations prevented the development of drug resistance

We hypothesized that since our predicted drug combinations target the main altered processes simultaneously, they may delay or prevent the development of drug resistance (**Fig. 7A**). To test this hypothesis, G361 and A375 cells were treated twice a week with single inhibitors or with different combinations of inhibitors, for 4-8 weeks.

**Figure 7:**
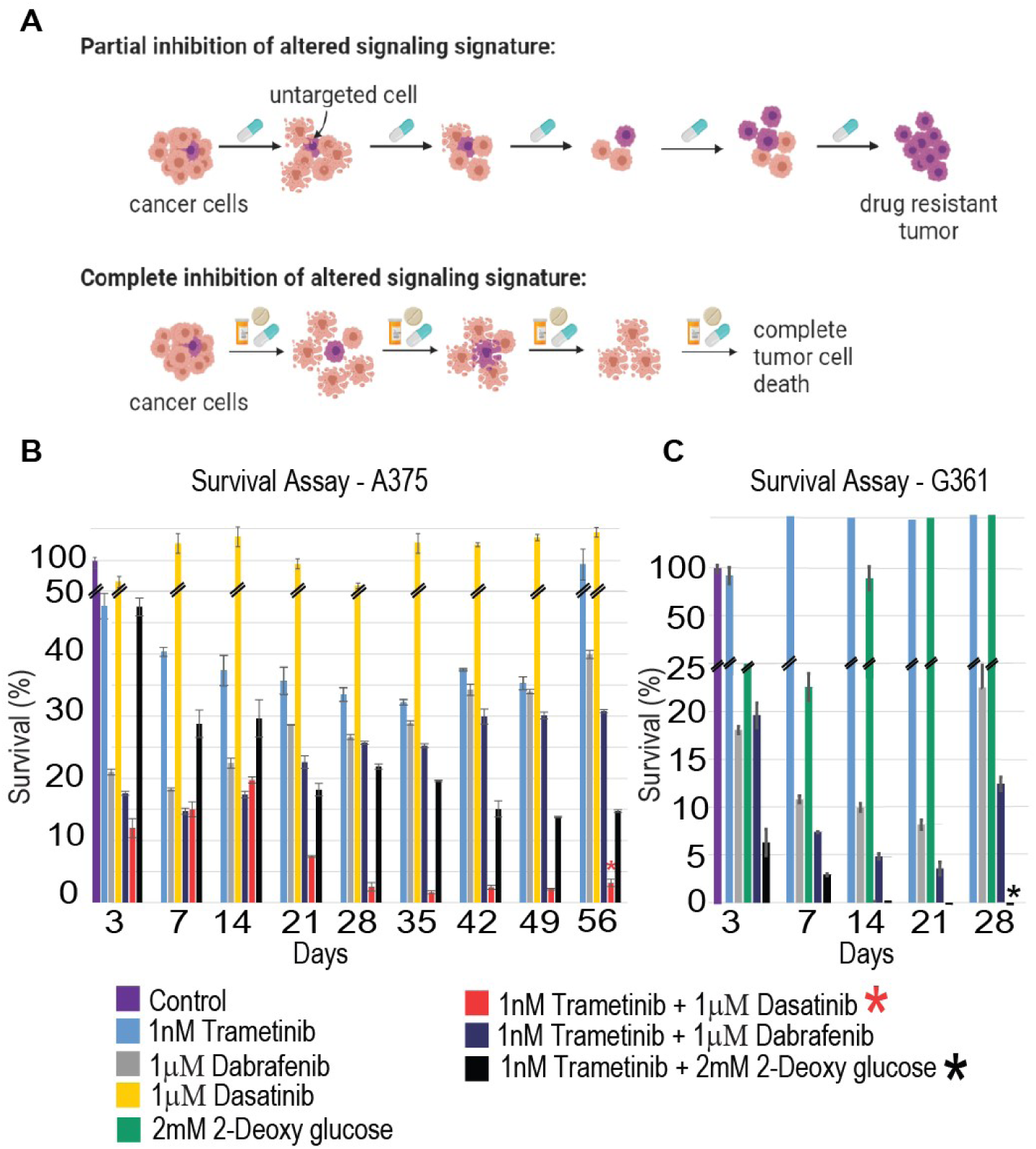
Development of resistance to different therapies. (**A**) The development of resistance to different types of therapies is shown in the illustration. The cells were treated with different therapies twice a week and then checked for cell survival. (**B**) A375 cells were treated with the monotherapies, trametinib+dasatinib, dabrafenib+trametinib or trametinib+2-deoxyglucose, twice weekly for 56 days. A development of resistance was evident after 21 days, but not in cells treated with trametinib+dasatinib. (**C**) G361 cells were treated with monotherapies, dabrafenib+trametinib, or trametinib+2-deoxyglucose, twice weekly for 28 days. The cells exhibited signs of drug resistance after 28 days. However, resistance development was not evident in cells that were treated with trametinb+2-deoxyglucose.

In G361 cells, 1 nM of trametinib demonstrated little to no effect on the survival of the cells (**Fig. 7B**). 1 µM of dabrafenib killed up to ∼92% of the cells at day 21, and then the cells began to regrow, even though the drug was still administered to the cells twice a week (**Fig. 7B**). 2 mM of 2-DG killed up to ∼78% of the cells at day 7, and then the cells began to regrow regardless of the presence of the drug (**Fig. 7B**). Combined treatment with trametinib and dabrafenib, a combination expected to partially target the altered signaling signature (**Fig. 5A**,**B**), effectively killed up to ∼96% of the cells at day 21, but then the cells began to regrow at day 28 in the presence of the drugs (**Fig. 7B**). However, when the cells were treated with the G361 PaSSS-based combination, trametinib and 2-DG (**Fig. 5A**,**B**), the cells continued to die until they reached a plateau at day 21, and no regrowth of the cells was evident (**Fig. 7B**).

Similar results were obtained in A375 cells: 1 nM trametinib killed 60% of the cells at day 7, and then the effect plateaued until the cells began to regrow at day 56 (**Fig. 7C**). 1 µM dabrafenib killed ∼78% of the cells at day 3, and then the cells kept growing till they reached 40% survival on day 56 (**Fig. 7C**). 1 µM dasatinib killed ∼40% of the cells at day 3, and then the cells regrew to 100% survival (**Fig. 7C**). Combined treatment with trametinib and dabrafenib achieved 88% killing at day 3, but then the cells grew until they plateaued at ∼30% survival at day 56 (**Fig. 7C**). Trametinib and 2-DG killed 55% of the cells at day 3 with an increase in effect over time, reaching a plateau of 15% survival at day 42 (**Fig. 7C**). The A375 PaSSS-based combination, trametinib and dasatinib (**Fig. 6A**,**B**), demonstrated a significant killing effect that became stronger with time, reaching near complete killing of the cells at 56 days (**Fig. 7C**).

These results clearly show that the PaSSS-based combinations predicted for each melanoma cell line prevent cellular regrowth in-vitro. Thus, targeting the actual altered signaling state, identified in the melanoma cells, and not necessarily the primary driver mutations, can be especially effective in disturbing the signaling flux and preventing cellular regrowth.

### The predicted drug combinations were superior to clinically used therapies in vivo

We turned to examine the effect of the PaSSS-predicted drug combination in murine models. The cells (A375 or G361) were injected subcutaneously into NSG mice, and treatments were initiated 6 times a week for up to 4 weeks (**Fig. 8**).

**Figure 8:**
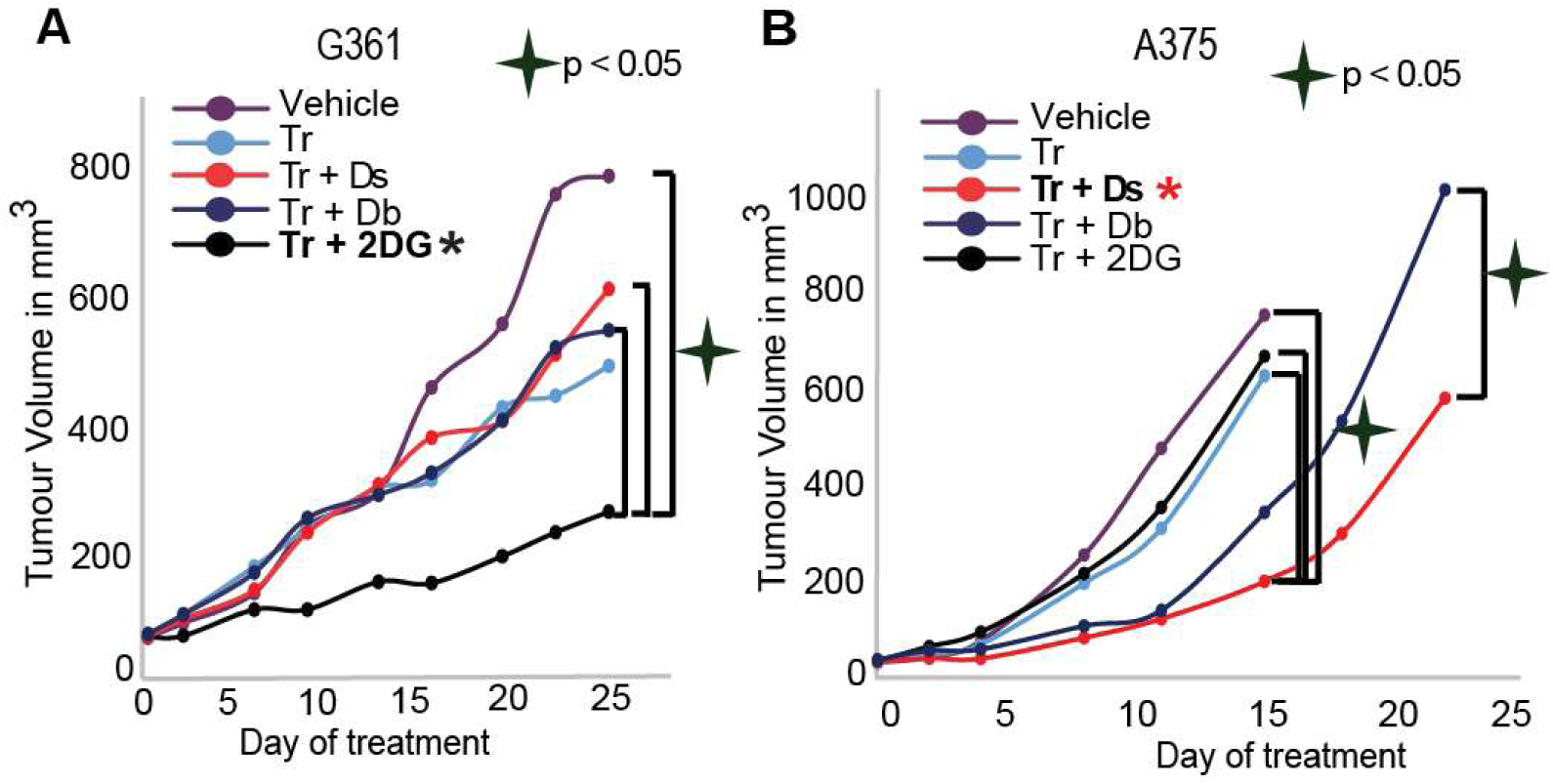
SA-based drug combinations demonstrated significantly reduced tumor growth *in vivo.* G361 or A375 (**B**) were injected subcutaneously into mice, and once tumors reached 50 mm^3^, treatments were initiated. In both cases, the PaSSS-based drug combinations, predicted to target the cell line-specific altered signaling signature, significantly inhibited tumor growth and demonstrated an effect superior to monotherapy of trametinib or to combinations predicted to partially target the PaSSS (see **Fig. 5**,**6** for details regarding the altered signaling signatures and PaSSS-based drug combination predictions).

Trametinib alone, or in combination with dasatinib or dabrafenib, was predicted to partially target the PaSSS of G361 cells (**Fig. 5A**,**B**). And indeed, these treatments demonstrated a reduction in tumor growth, relative to vehicle treatment (**Fig. 8A**). However, the PaSSS-based combination, trametinib + 2-DG, demonstrated the strongest effect, and achieved significant inhibition of G361 tumor growth (**Fig. 8A**).

A375 tumors that were treated with trametinib alone or with a combination trametinib + 2-deoxyglucose (predicted to be efficient for G361 but not for A375 cells (**Fig. 5, 6**)) demonstrated slightly reduced growth relative to vehicle-treated tumors (**Fig. 8B**). When A375 tumors were treated with the clinically used combination, trametinib + dabrafenib, a stronger effect was observed (**Fig. 8B**). PaSSS analysis predicted that trametinib + dabrafenib would achieve partial inhibition of the altered signaling in A375 cells (**Fig. 6A**,**B**), and that adding dasatinib to trametinib should achieve inhibition of intracellular signaling that have emerged in A375 cells (**Fig. 6A**,**B**). Indeed, the combination trametinib + dasatinib demonstrated an effect superior to all other treatments, and significantly inhibited the growth of A375 tumors (**Fig. 8B**).

These results point to a significantly higher efficiency of the PaSSS-predicted combinations relative to drug combinations used in clinics. Moreover, we demonstrated the selectivity of the individualized treatments. The predicted and very effective combination for one BRAF^V600E^ melanoma malignancy was significantly less effective for the other, and vice versa (**Fig. 8**). Our results underscore the need for a personalized treatment for each melanoma patient.

## Discussion

With the accelerated gain of knowledge in the field of melanoma therapy and cancer research, it is becoming clear that tumors evolving from the same anatomical origins cannot necessarily be treated the same way [40]. Inter tumor heterogeneity results in various response rates of patients to therapy [41–43]. Herein we extend this notion, and show that even tumors that were initially driven by the same oncogenes, specifically BRAF^V600E^-driven melanoma tumors, often evolve in different molecular manners [44], giving rise to distinct altered signaling signatures, or PaSSS (patient-specific altered signaling signature), at the time of biopsy.

We show that 17 altered molecular processes are repetitive among the 725 SKCM and THCA tumors. Each tumor is characterized by a specific PaSSS, i.e. a subset of ∼1-3 unbalanced processes. Accordingly, each patient is assigned a unique barcode, denoting the normalized PaSSS. We show that the collection of 725 tumors is described by 138 distinct barcodes, suggesting that the cohort of patients consists of 138 types of cancer, rather than only 4 types (SKCM or THCA; BRAF^WT^ or BRAF^V600E^). These 138 types of tumors, each representing a barcode, or a sub-combination of 17 unbalanced processes, are mapped into a multi-dimensional space, consisting of 17 dimensions. Once the tumor-specific information is transformed into a multi-dimensional space, treating these thousands of tumors becomes at an arm’s reach. The specific barcode assigned to each patient allows the rational design of patient-tailored combinations of drugs, many of which already exist in clinics.

We found that 353 BRAF^V600E^ and BRAF^WT^ melanoma tumors are described by 87 distinct barcodes of unbalanced processes, and that 372 BRAF^V600E^ and BRAF^WT^ THCA tumors are described by 51 barcodes. Interestingly, the barcodes appeared to be almost mutually exclusive between SKCM and THCA tumors (**Table S4**). While this finding suggests that the molecular processes underlying SKCM and THCA tumor evolution may have organ-specific differences, the large number of cancer type-specific barcodes and the large number of barcodes describing single patients underscore the need for personalized diagnosis and treatment.

We show that tumors harboring BRAF^V600E^ can harbor distinct PaSSSs, and in contrast, that tumors can harbor the same PaSSS regardless of whether they carry BRAF^V600E^ or BRAF^WT^. We therefore deduce that profiling melanoma patients according to their BRAF mutational status is insufficient to assign effective therapy to the patient. Since the unbalanced processes each harbor a specific group of co-expressed altered proteins, they should all be targeted simultaneously to reduce the altered signaling flux in the tumor.

We demonstrate this concept experimentally by analyzing a cell line dataset and predicting efficient targeted drug combinations for two selected BRAF^V600E^ melanoma cell lines, G361 and A375. We show that although both cell lines contain the mutated BRAF^V600E^, they harbor distinct barcodes, and demand different combinations of drugs (**Fig. 5, 6**). We demonstrate that in both cell lines, our PaSSS-based combinations indeed achieved efficient killing of the cells and reached a killing rate that was higher than that of the drug combination often prescribed clinically to BRAF^V600E^ patients, dabrafenib+trametinib. These results were recapitulated in vivo as well (**Fig. 8**). Moreover, we demonstrated the selectivity of the PaSSS-based drug combinations. The highly efficient PaSSS-based drug combination for one melanoma malignancy can be significantly less efficient for another melanoma and vice versa.

The results reported here highlight the urgent need for the design of personalized treatments for melanoma patients based on individualized alterations in signaling networks rather than on initial mutational events. Furthermore, the study establishes PaSSS analysis as an effective approach for the design of personalized cocktails comprising FDA-approved drugs. Personalized targeted cocktails, which may be further combined with immunotherapy strategies, are expected to provide long term efficacy for melanoma patients.

## Acknowledgements

The funding sources for this work were from Israel Science Foundation (ISF) and NIH.

## Availability of data and materials

The human tumor dataset that supports the findings of this study is publicly available for download from the TCPA portal [14], https://tcpaportal.org/tcpa/download.html > Pan-Can 32.

The cell line dataset that supports the findings of this study is publicly available for download from the TCPA portal [14], https://tcpaportal.org/mclp/#/datasets

## Authors’ contributions

N.K.B, S.V. and E.F.A designed the research, N.K.B, E.F.A and S.V carried out computational analyses, E.F.A, S.V, I.A.A, D.V., S.S., and A.A. performed experiments, N.K.B, S.V., and E.F.A wrote the manuscript. All authors approved the manuscript.

## Consent for publication

The authors confirm that this manuscript does not contain any personal data or images from any individual participants.

