## Supplementary material for "Overcoming Resistance to BRAF^V600E^ Inhibition in Melanoma by Deciphering and Targeting Personalized Protein Network Alterations": SI

Swetha Vasudevan[#], Efrat Flashner-Abramson[#], Ibukun Adesoji Adejumobi, Dana Vilencki, Shira Stefansky, Ariel Rubinstein and Nataly Kravchenko-Balasha*

Department for Bio-medical Research, Faculty of Dental Medicine, Hebrew University of Jerusalem, Jerusalem 91120, Israel

### Equal contribution

### Supplementary Figures

#### Figure S1.

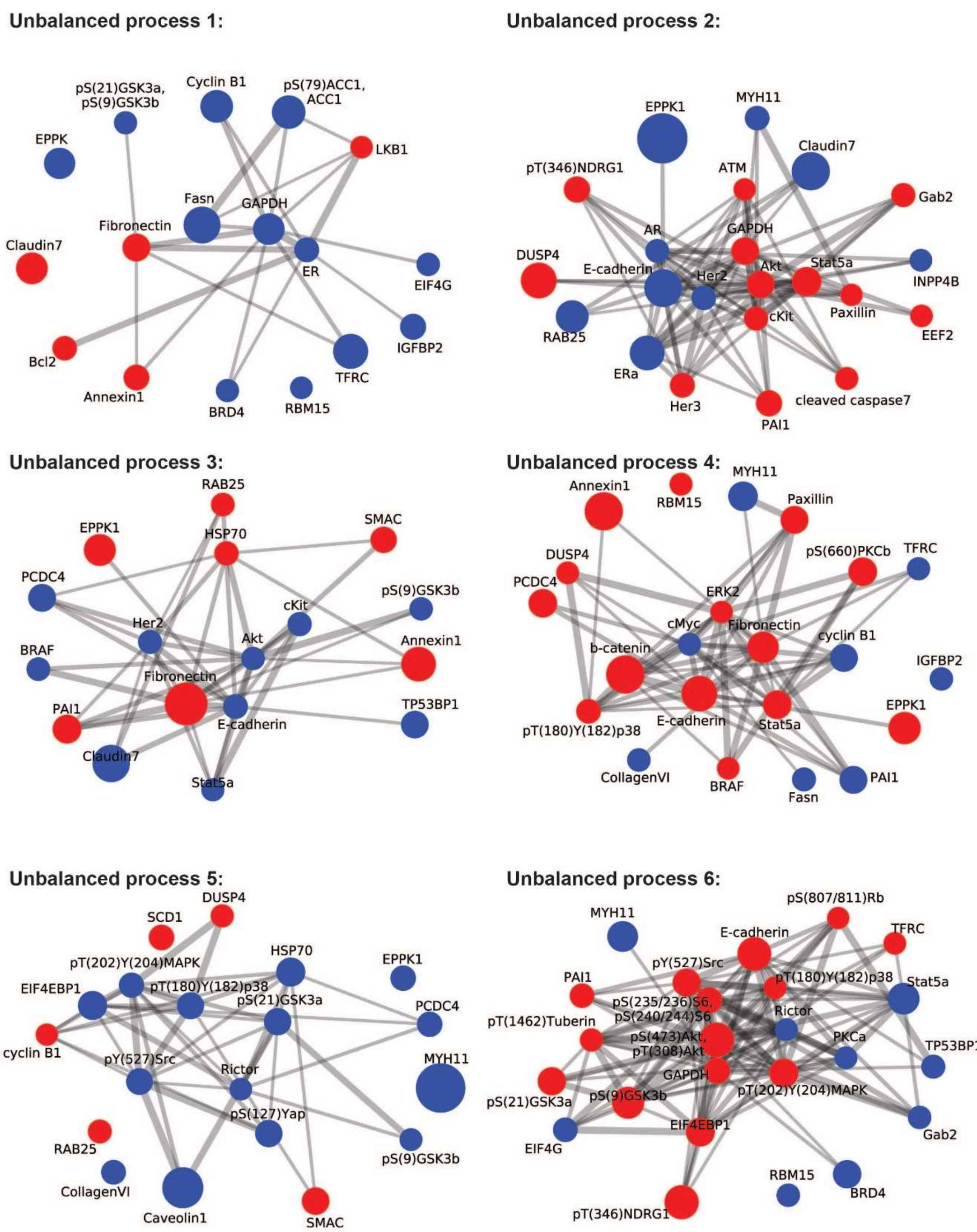

**Unbalanced process 7:**

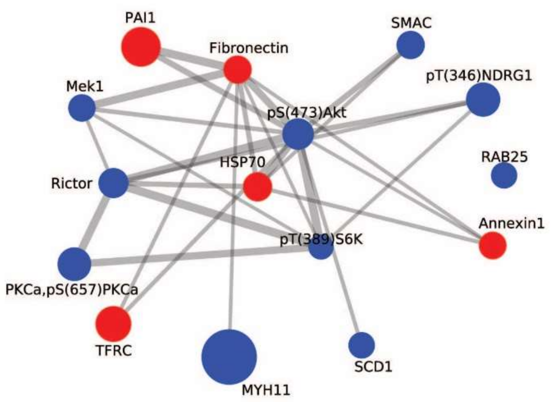

**Unbalanced process 8:**

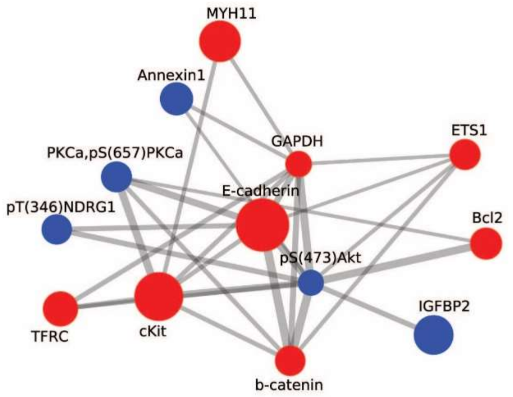

**Unbalanced process 9:**

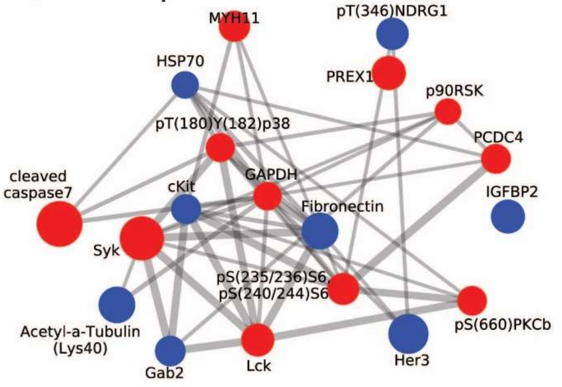

**Unbalanced process 10:**

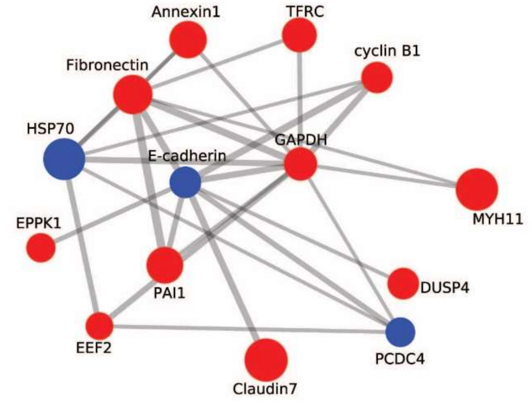

**Unbalanced process 11:**

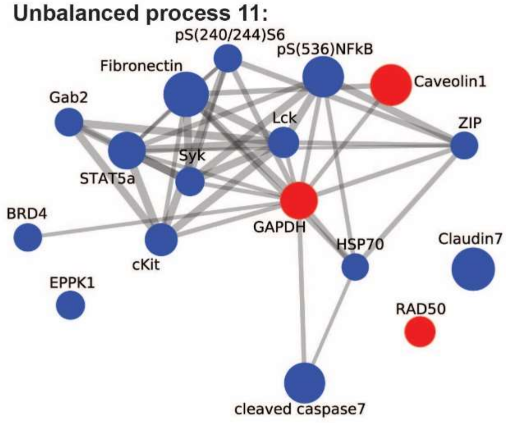

**Unbalanced process 12:**

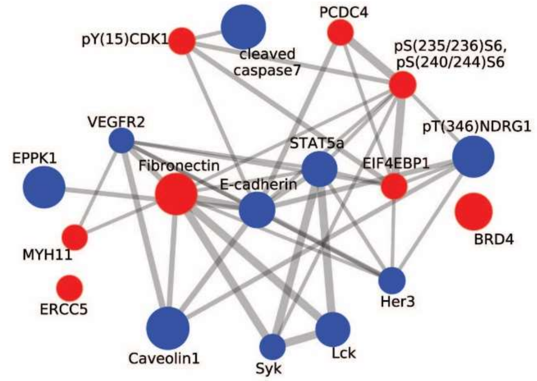

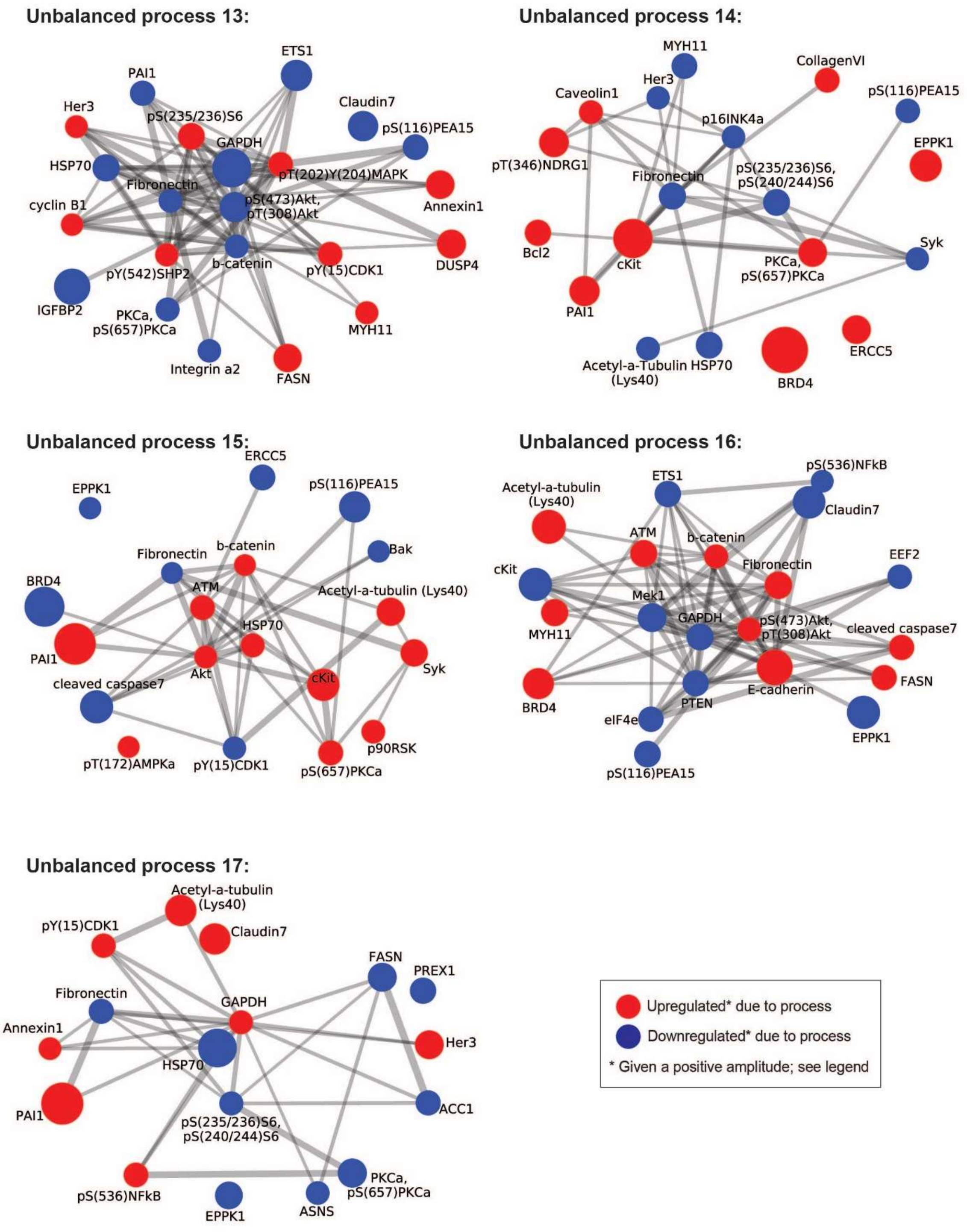

**Figure.S1 The unbalanced subnetworks identified by PaSSS analysis for 725 SKCM and THCA tumors.** For every process α, the proteins were assembled into networks using functional interactions according to STRING database. **<u>Note</u>** that red proteins are upregulated, and blue proteins are downregulated <u>given that the amplitude of the process is positive</u>. In tumors where the amplitude is negative, the direction of change is opposite.

**Figure S2.**

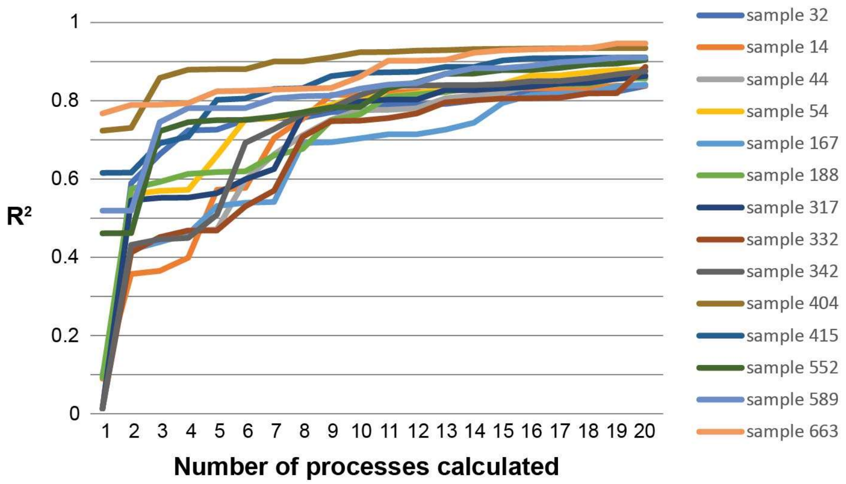

**Figure S2. 17 unbalanced processes suffice to describe the biological imbalance in 725 tumors.** Presented are $R^2$ values for 14 random patients. The $R^2$ was calculated for every patient by plotting the natural logarithm of the experimental data (LnXi) vs. $\sum G_{i\alpha}\lambda_{\alpha}(k)$ for $\alpha = 0,1,2,\ldots,20$ . As more constraints are added, we expect to get higher agreement between the experimental data and the theoretical calculation, reaching the highest level of correlation when the experimental data is fully reproduced by the theoretical fitting. The plot reaches a plateau after the $17^{th}$ constraint. This indicates that for $\alpha = 18,19,20\ldots$ the data includes mainly noise.

**Figure S3.**

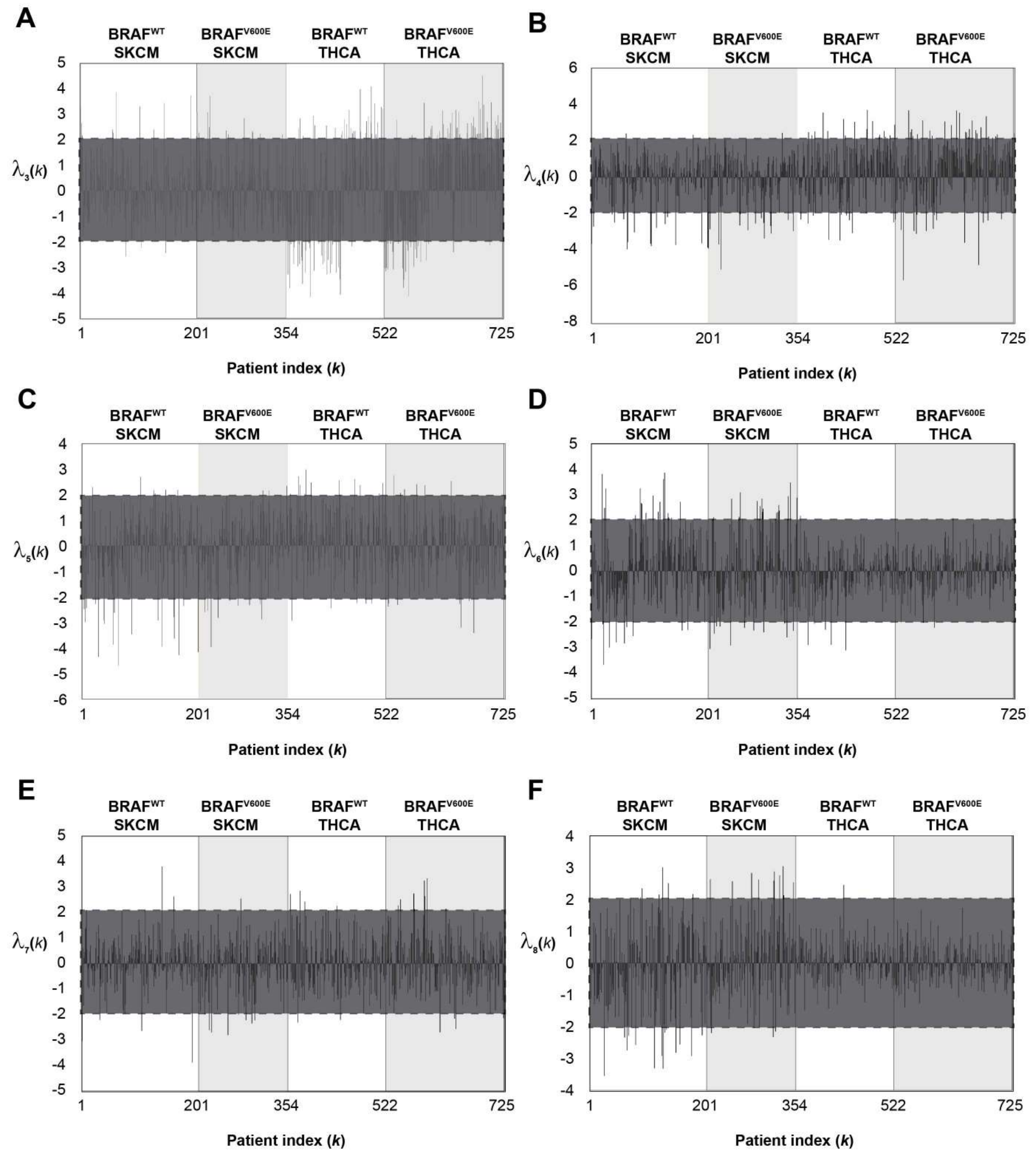
A
B
C
D
E
F
BRAF^WT SKCM
BRAF^V600E SKCM
BRAF^WT THCA
BRAF^V600E THCA
λ3(k)
λ4(k)
λ5(k)
λ6(k)
λ7(k)
λ8(k)
1
201
354
522
725
Patient index (k)

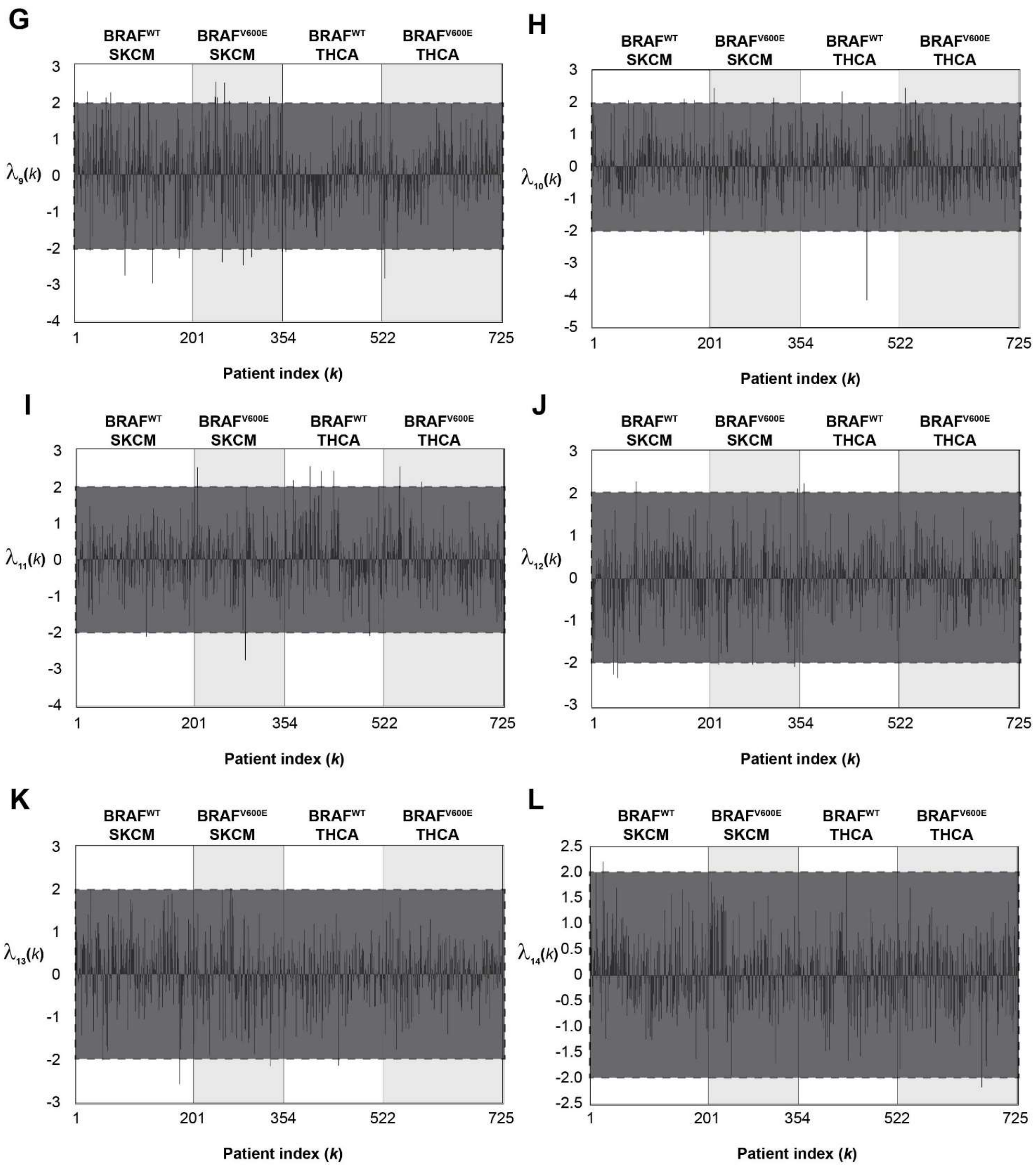
G
BRAF WT SKCM
BRAF V600E SKCM
BRAF WT THCA
BRAF V600E THCA
λ9(k)
Patient index (k)
1 201 354 522 725
H
BRAF WT SKCM
BRAF V600E SKCM
BRAF WT THCA
BRAF V600E THCA
λ10(k)
Patient index (k)
I
BRAF WT SKCM
BRAF V600E SKCM
BRAF WT THCA
BRAF V600E THCA
λ11(k)
Patient index (k)
J
BRAF WT SKCM
BRAF V600E SKCM
BRAF WT THCA
BRAF V600E THCA
λ12(k)
Patient index (k)
K
BRAF WT SKCM
BRAF V600E SKCM
BRAF WT THCA
BRAF V600E THCA
λ13(k)
Patient index (k)
L
BRAF WT SKCM
BRAF V600E SKCM
BRAF WT THCA
BRAF V600E THCA
λ14(k)
Patient index (k)

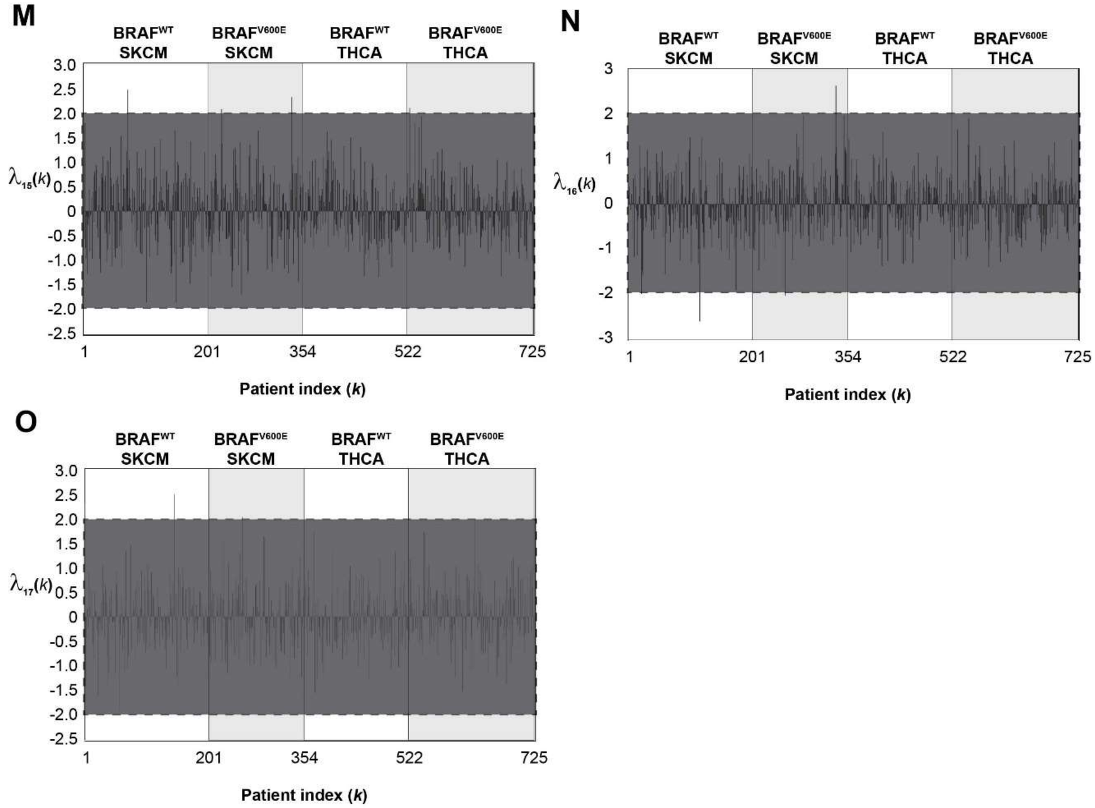

**Figure S3. Tumor-specific amplitudes ($\lambda_\alpha(k)$) of the 17 unbalanced processes.** Not all unbalanced processes are active in all patients. Rather, each unbalanced process, $\alpha$, is assigned an amplitude, $\lambda_\alpha(k)$, for the specific tumor, $k$ (see also **Table S1**). Every panel in the figure presents 725 values of $\lambda_\alpha(k)$ for a specific value of $\alpha$. The gray boxes mark the threshold limits – only values greater than 2 or smaller than -2 were considered significant (see Methods for more details).

**Figure S4.**

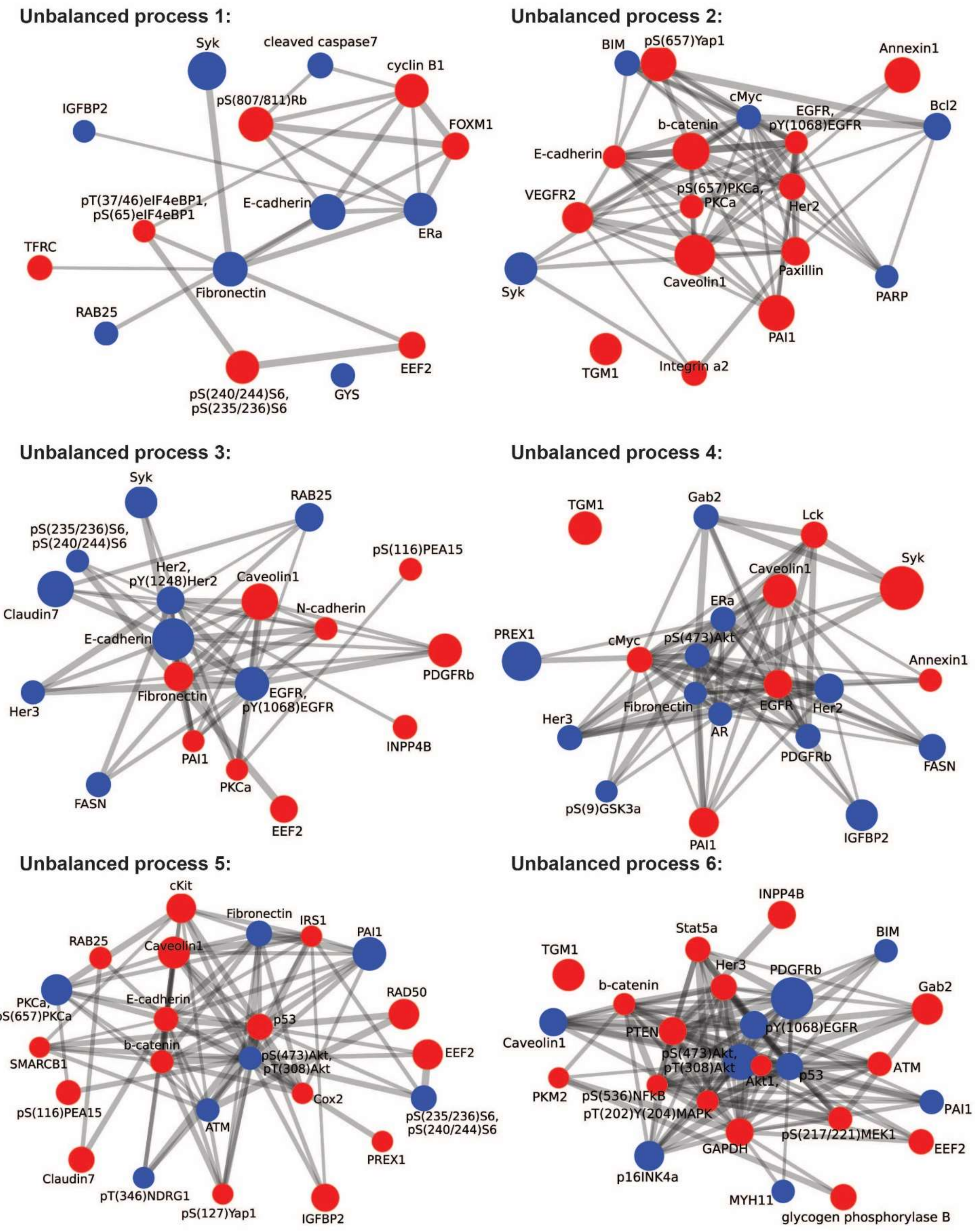
Unbalanced process 1:
Syk
cleaved caspase7
cyclin B1
IGFBP2
pS(807/811)Rb
FOXM1
pT(37/46)eIF4eBP1,
pS(65)eIF4eBP1
E-cadherin
ERa
TFRC
Fibronectin
RAB25
EEF2
pS(240/244)S6,
pS(235/236)S6
GYS
Unbalanced process 2:
BIM
pS(657)Yap1
Annexin1
cMyc
Bcl2
b-catenin
EGFR,
pY(1068)EGFR
E-cadherin
pS(657)PKCa,
PKCa
VEGFR2
Her2
Syk
Caveolin1
Paxillin
PARP
PAI1
TGM1
Integrin a2
Unbalanced process 3:
Syk
RAB25
pS(235/236)S6,
pS(240/244)S6
pS(116)PEA15
Her2,
pY(1248)Her2
Caveolin1
Claudin7
N-cadherin
E-cadherin
PDGFRb
Fibronectin
EGFR,
pY(1068)EGFR
Her3
INPP4B
PAI1
PKCa
FASN
EEF2
Unbalanced process 4:
TGM1
Gab2
Lck
Syk
Caveolin1
ERa
PREX1
cMyc
pS(473)Akt
Annexin1
Fibronectin
EGFR
Her2
Her3
AR
PDGFRb
FASN
pS(9)GSK3a
PAI1
IGFBP2
Unbalanced process 5:
cKit
Fibronectin
IRS1
RAB25
Caveolin1
PAI1
RAD50
PKCa,
pS(657)PKCa
E-cadherin
p53
b-catenin
pS(473)Akt,
pT(308)Akt
EEF2
SMARCB1
Cox2
pS(116)PEA15
ATM
pS(235/236)S6,
pS(240/244)S6
PREX1
Claudin7
pT(346)NDRG1
pS(127)Yap1
IGFBP2
Unbalanced process 6:
INPP4B
Stat5a
TGM1
BIM
Her3
PDGFRb
b-catenin
Gab2
PTEN
pY(1068)EGFR
Caveolin1
pS(473)Akt,
pT(308)Akt
Akt1,
p53
ATM
PKM2
pS(536)NFkB
PAI1
pT(202)Y(204)MAPK
pS(217/221)MEK1
GAPDH
EEF2
p16INK4a
MYH11
glycogen phosphorylase B

**Unbalanced process 7:**

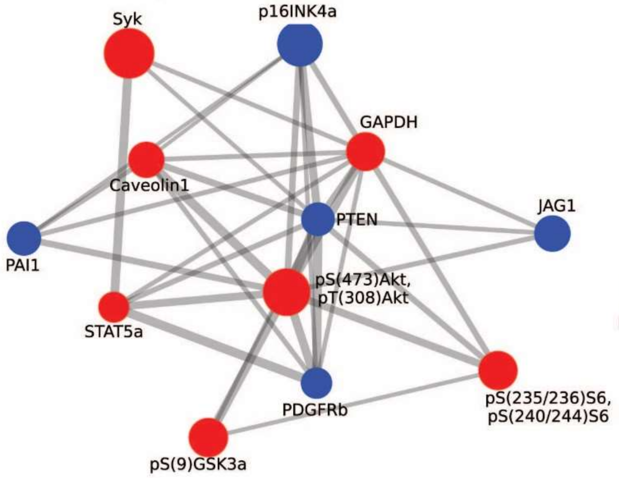

**Unbalanced process 8:**

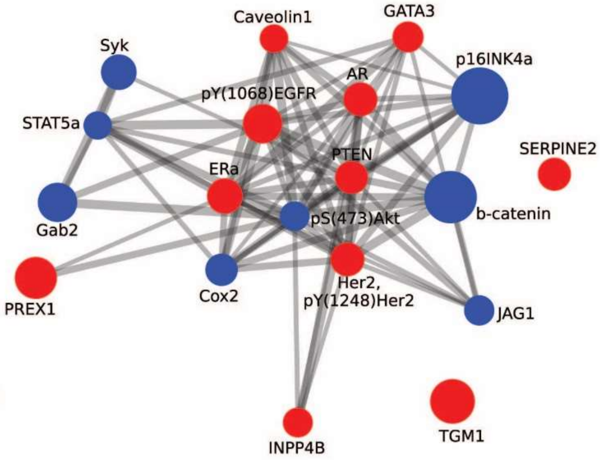

**Unbalanced process 9:**

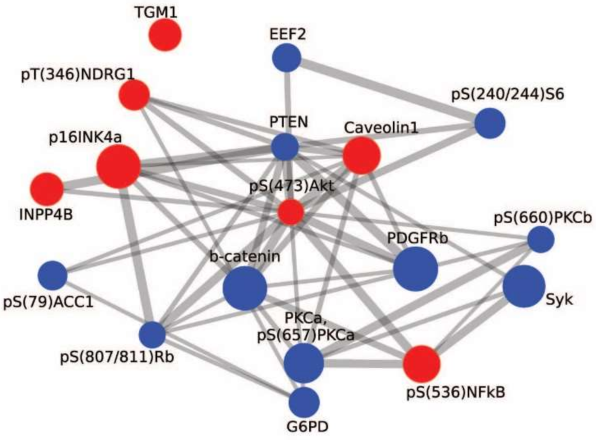

**Unbalanced process 10:**

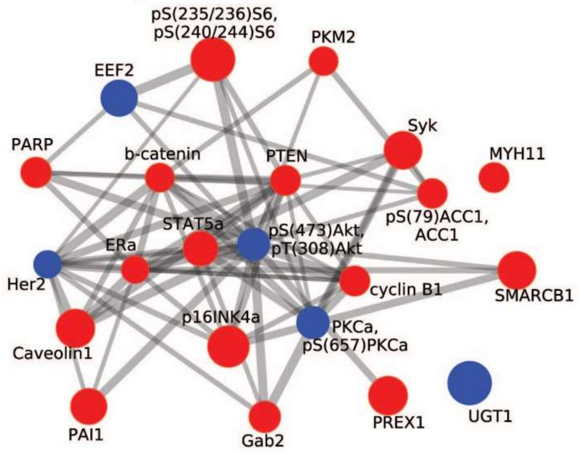

**Unbalanced process 11:**

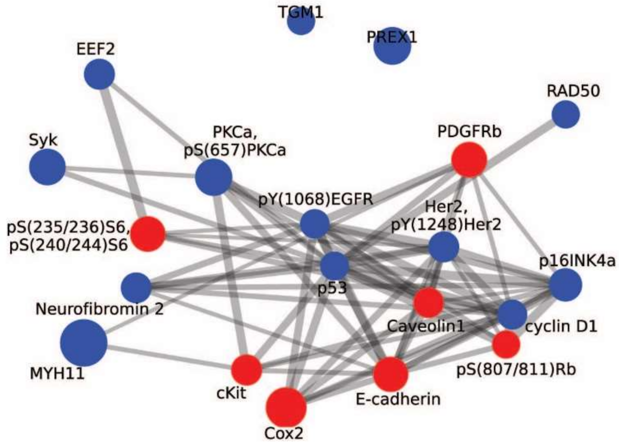

**Unbalanced process 12:**

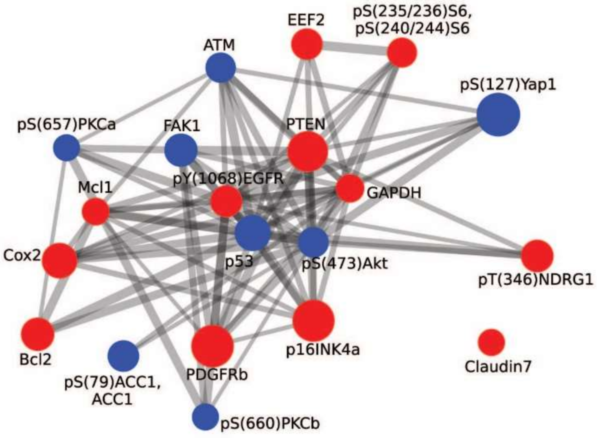

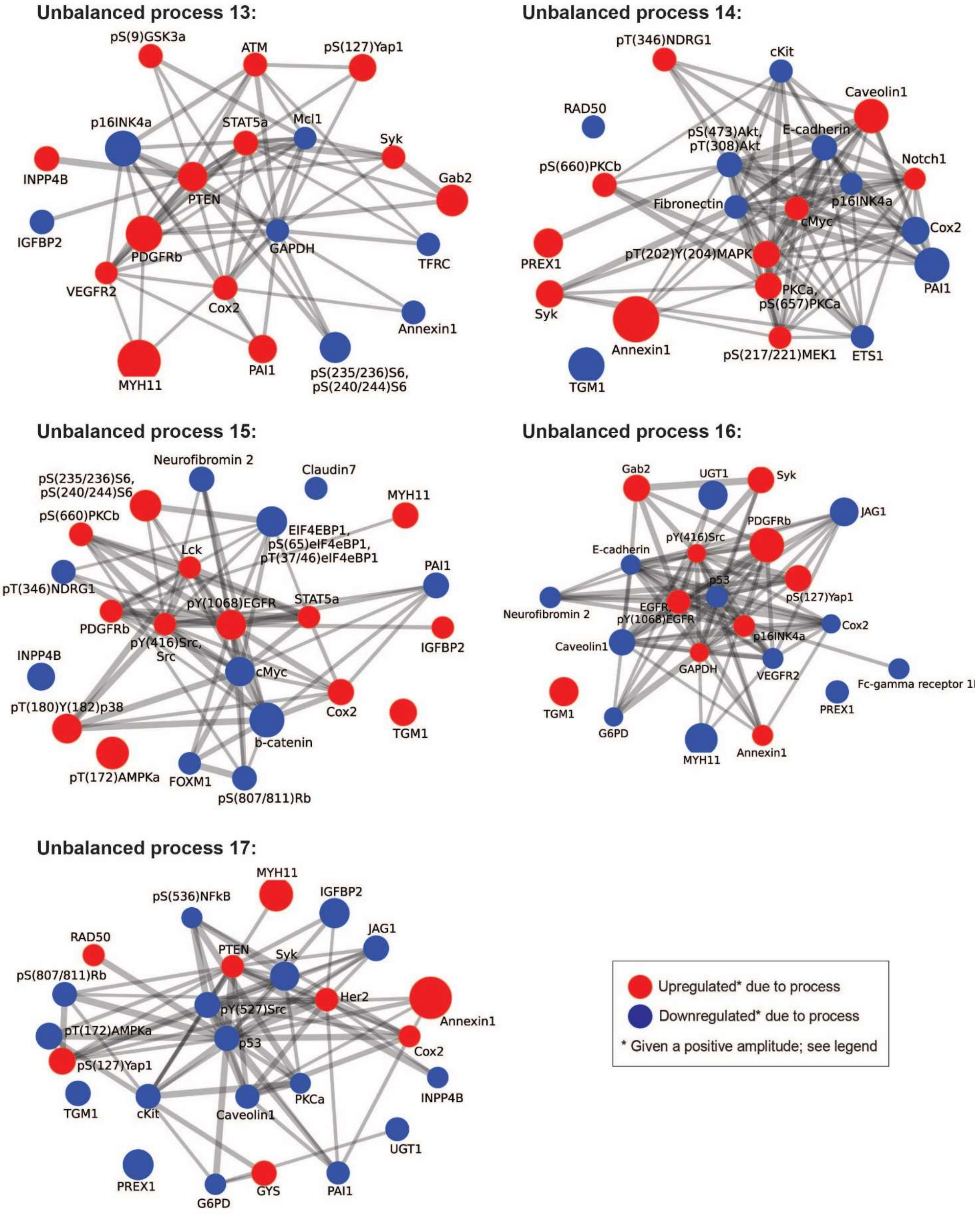

**Figure S4. The unbalanced subnetworks identified by PaSSS analysis for 219 cell lines.** For every process α, the proteins were assembled into networks using functional interactions according to STRING database. **Note** that red proteins are upregulated, and blue proteins are downregulated given that the amplitude of the process is positive. In tumors where the amplitude is negative, the direction of change is opposite.
